# Transcriptome analysis provides genome annotation and expression profiles in the central nervous system of *Lymnaea stagnalis* at different ages

**DOI:** 10.1101/2021.05.17.444496

**Authors:** Martina Rosato, Brittany Hoelscher, Zhenguo Lin, Chidera Agwu, Fenglian Xu

## Abstract

Molecular studies of the freshwater snail *Lymnaea stagnalis*, a unique model organism for neurobiology research, has been severely hindered by the lack of sufficient genomic information. As part of our ongoing effort studying *L. stagnalis* neuronal growth and connectivity at various developmental stages, we provide the first age-specific transcriptome analysis and gene annotation of young, adult, and old *L. stagnalis* central nervous system (CNS). RNA sequencing using Illumina NovaSeq 6000 platform produced 56-69 millions of 150 bp paired-end reads, and 74% of these reads were mapped to the draft genome of *L. stagnalis*. We provide gene annotations for 32,288 coding sequences with a minimum of 100 codons, contributing to the largest number of annotated genes for the *L. stagnalis* genome to date. Lastly, transcriptomic analyses reveal age-specific differentially expressed genes and enriched pathways in young, adult, and old CNS. These datasets represent the largest and most updated *L. stagnalis* CNS transcriptomes.

## Introduction

The freshwater snail *Lymnaea stagnalis* belongs to the phylum Mollusca, class Gastropoda (Kuroda & Abe, 2020). Like its counterpart, the sea slug *Aplysia californica, L. stagnalis* has served as an important mollusc model organism for the neurobiology field since the 1970s due to its simple central nervous system (CNS) (Fodor, Hussein, Benjamin, Koene, & Pirger, 2020). *L. stagnalis* CNS contains a total of 20,000-25,000 neurons organized in a ring of 11 connected ganglia. The neurons are large in size (up to ∼100 µm in diameter) and easily recognizable, making them a perfect target for *in vitro* and *in vivo* studies. Many studies have used this model to investigate the fundamental mechanisms of neuronal networks involved in various behaviours including feeding (Kojima, Nanakamura, Nagayama, Fujito, & Ito, 1997; Yeoman, Kemenes, Benjamin, & Elliott, 1994), respiration (Haque et al., 2006; Taylor & Lukowiak, 2000), locomotion (Syed & Winlow, 1991; Vorontsov, Tsyganov, & Sakharov, 2004), and reproduction (Hermann, de Lange, Pieneman, ter Maat, & Jansen, 1997; Jimenez et al., 2004). Studies have also focused on high cognitive behaviours, including learning and memory (Dodd, Rothwell, & Lukowiak, 2018; Sunada et al., 2017; Swinton et al., 2019; Tan & Lukowiak, 2018), as well as deciphering cellular mechanisms of synapse formation and synaptic plasticity during development (Getz, Wijdenes, Riaz, & Syed, 2018; Mersman, Jolly, Lin, & Xu, 2020; Onizuka et al., 2012). *L. stagnalis* has also recently gained increasing popularity for the investigation of brain aging and neurodegenerative diseases such as Parkinson’s disease and Alzheimer’s disease (Arundell et al., 2006; de Weerd, Hermann, & Wildering, 2017; Ford, Crossley, Vadukul, Kemenes, & Serpell, 2017; Hermann, Perry, Hamad, & Wildering, 2020; Maasz et al., 2017). It is important to note that comparative studies have highlighted several human homologs involved in aging and neurodegenerative disease in both *A. californica* and *L. stagnalis* (Fodor, Urban, Kemenes, Koene, & Pirger, 2020; Moroz et al., 2006; Moroz & Kohn, 2010), showing the great potential for future molecular insights into brain aging and pathology using these unique mollusc models. More importantly, a recent study has successfully established the use of CRISPR/Cas9 in *L. stagnalis* embryos (Abe & Kuroda, 2019), further underscoring the high feasibility of *L. stagnalis* for genetic studies.

Despite the importance of *L. stagnalis* to brain network, behaviour, and development studies, genetic information is mostly limited to the identification and cloning of individual genes. Only in the past decade have large-scale genomic analyses been put forward to characterize the *L. stagnalis* transcriptome. For example, several studies (Bouetard et al., 2012; Davison & Blaxter, 2005; Feng et al., 2009) have provided transcript sequencing data using expressed sequence tags (EST) generated from *L. stagnalis* CNS libraries. Although these data have provided valuable insights into partial gene expression, they were insufficient to perform transcriptome analysis due to the limitation of the EST-based technique. A later study by Sadamoto et al. in 2012 (Sadamoto et al., 2012) took advantage of the development of deep RNA sequencing (RNA-Seq) techniques and performed de novo transcriptome shotgun assembly (TSA) on *L. stagnalis* RNA samples of CNS. This study provided improved transcriptome data with longer and larger sequences and also contributed to the identification of novel transcripts in *L. stagnalis* CNS compared to previous studies. Moreover, from both Blast searches in public databases and comparison with previous *L. stagnalis* and *A. californica* EST, they showed that very few of their sequences had blast hits. This result was mainly attributed to the lack of sufficient molluscan sequence coverage in the public databases for valid dataset comparison, urging the need for continued genetic studies of *L. stagnalis* and other gastropods. A very recent effort by Dong et al. (Dong et al., 2021) has established an updated *L. stagnalis* transcriptome of adult CNS by RNA-Seq. However, the above studies have focused on only one developmental time point, predominantly in adults. Considering the increasing use of *L. stagnalis* for brain aging and pathology research (Fodor, Urban, et al., 2020), updated transcriptome datasets and gene annotations including old or aging snail CNS are critically needed.

Brain development, maturation, and aging are influenced by both intrinsic (genetic) and extrinsic (environmental) factors throughout the life span of animals and human. Large-scale study of transcriptional changes in brains of animals at various ages provide important molecular insights into brain development, aging, pathology, and evolution. Spatial and/or temporal transcriptome analyses of brains and other tissues have been carried out in human (Kang et al., 2011; Tebbenkamp, Willsey, State, & Sestan, 2014), rats (Shavlakadze et al., 2019), mice (Chou et al., 2016), chicken (Xu et al., 2018), zebrafish (Vesterlund, Jiao, Unneberg, Hovatta, & Kere, 2011), and birds (Frias-Soler, Pildain, Parau, Wink, & Bairlein, 2020) among others. All these studies have contributed to our understanding of the molecular basis of brain development. Invertebrates have also been utilized for study of development and aging. Developmental transcriptomes of well-established invertebrate models such as *Caenorhabditis elegans* (*C. elegans*) (Boeck et al., 2016; Lu, Lai, Liao, & Tsai, 2020) and *Drosophila melanogaster* (*D. melanogaster*) (Graveley et al., 2011) have been reported. Recent efforts have also focused on the transcriptome of aging *D. melanogaster* (Moskalev et al., 2019; Pacifico, MacMullen, Walkinshaw, Zhang, & Davis, 2018) and *C. elegans* (Tarkhov et al., 2019), aiming to reveal molecular mechanisms of longevity or aging trajectories.

In mollusca, the developmental (embryonic, larval, and metamorphic) transcriptome of *A. californica (Heyland, Vue, Voolstra, Medina, & Moroz, 2011)* and maternal (1 to 2 cell and ∼32 cell) transcriptome of *L. stagnalis* have been conducted (Liu, Davey, Jackson, Blaxter, & Davison, 2014). These studies shed novel insights into conserved sets of genes and pathways in early development. However, these studies failed to inform how these genes or other sets of genes are regulated in later stages of life, such as after animals are fully matured and aged. Although whole-transcriptome changes in tail-withdrawal sensory neurons of sexually matured and aged *A. californica* have been reported (Greer, Schmale, & Fieber, 2018), transcriptome changes of entire CNS in young, mature, and aged *A. californica* and *L. stagnalis* have not been carried out. Such studies are critical for our complete understanding and comparative studies of age- or species-specific molecular strategies that are key to the evolution, survival, and function of both invertebrate and vertebrate.

To this end, in the present study, we provide whole transcriptome analysis in *L. stagnalis* CNS from three different ages: 3 months (young), 6 months (adult), and 18 months (old). This is the first time that changes in CNS transcriptome profiling during brain development, maturation, and aging in *L. stagnalis* are analyzed. *L. stagnalis* has a relatively short life cycle, with a life expectancy of about 1.5 to 3 years (Hermann et al., 2007). The embryonic stage of the snail lasts for around two weeks, and eggs are contained in gelatinous masses that are accessible for genetic manipulation. After hatching, young snails reach sexual maturity at around 4 to 6 months of age, and senescence starts after 7-8 months (Hermann et al., 2007; Janse, Slob, Popelier, & Vogelaar, 1988). Therefore, the 3-month-old age in our study represents a rapid developing and sexual immature stage, the 6-month-old age represents a fully, sexually mature stage, and 18-month-old represents an aging stage (Hermann et al., 2007; Janse et al., 1988).

Using the above three age cohorts, our study generated 55-69 millions of 150 bp paired-end RNA-Seq reads using the Illumina NovaSeq 6000 platform. Of these reads, ∼74% were successfully mapped to the unannotated reference genome of *L. stagnalis*. Our reference-based transcriptome assembly yielded 42,478 gene loci, of which 37,661 genes encode coding sequences (CDS) of at least 100 codons. In addition, we provide gene annotations for 32,288 out of 37,661 (∼88%) of these sequences, contributing the largest number of annotated genes in *L. stagnalis* CNS so far. Moreover, among 242 previously cloned *L. stagnalis* genes, we were able to match ∼87% of them in our transcriptome, a high percentage of gene coverage. The changes in gene transcription levels were validated by real-time qPCR for three *innexin* genes: *Inx1, Inx4* and *Inx5*. Lastly, our transcriptomic analyses revealed distinct, age-specific gene clusters, differentially expressed genes, and enriched pathways in young, adult, and old CNS. Together, these datasets are the largest and most updated *L. stagnalis* CNS transcriptomes, which will serve as a resource for future molecular studies and functional annotation of transcripts and genes in *L. stagnalis*.

## Materials and Methods

### Animals and brain dissection

*L. stagnalis* were maintained in artificial pond water at 20°C in a 12-hour light/dark cycle and fed with romaine lettuce twice a week. *L. stagnalis* were obtained from the University of Calgary, Canada (original stock was from the Vrije University in Amsterdam) and raised and maintained in aquaria at Saint Louis University since 2015 according to protocols developed and optimized as described previously (Mersman et al., 2020; Steen, Jager, & Hoven, 1968). All procedures are in accordance with the standard operating protocol guidelines established by the U.S. Department of Agriculture Animal and Plant Health Inspection Service. Animals at 3 months old (young), 6 months old (adult) or 18 months old (old) were used for RNA sequencing and qPCR. We used replicate samples for each developmental age, and for each sample, the CNS of ten animals were pooled. The snails were de-shelled and anesthetized in 10% (v/v) Listerine in *L. stagnalis* saline (51.3 mM NaCl; 1.7 mM KCl; 4.0 mM CaCl_2_; 1.5 mM MgCl_2_, 10 mM HEPES, pH 7.9), and the dissected central ring ganglia were used for both RNA sequencing and qPCR.

### RNA extraction

RNA was extracted from dissected *L. stagnalis* central ring ganglia using the RNeasy Mini Kit (Qiagen, 74104) following the manufacturer’s instructions. RNA concentration was assessed using a Nanodrop 2000 Spectrophotometer (ThermoFisher, ND-2000). After RNA extraction, genomic DNA (gDNA) was removed via the TURBO DNA-free kit (Invitrogen, AM1907).

### RNA sequencing library construction, sequencing, alignment, and transcriptome assembly

The construction of RNA sequencing libraries using polyA enrichment method was performed by Novogene Corporation Inc (Sacramento, CA, USA). These libraries were sequenced using the Illumina NovaSeq 6000 platform (Paired-end, 150 bp, insert size 250-300 bp). The sequencing reads of each RNA-Seq library were aligned to the reference genome of *L. stagnalis* (assembly v1.0 GCA_900036025.1) using HISAT2 (Kim, Langmead, & Salzberg, 2015). The soft clipping option in HISAT2 was enabled to exclude low-quality bases at both ends of reads. The number of reads in each sample successfully mapped to the *L. stagnalis* reference genome are provided in **Supplemental table 1**. We used Stringtie (Pertea et al., 2015) to assemble transcripts based on aligned reads. The expression abundance of each transcript was quantified as fragment per kilobase million reads (FPKM). The results of principal component analyses (PCA) of the abundance of transcripts (FPKM) for all genes from all samples are provided in **Supplemental Fig. 1A**. The correlations of gene expression profile between each pair of samples are provided in **Supplemental Fig. 1B**. The raw sequencing data generated in this study have been submitted to the NCBI BioProject database under accession number PRJNA698985.

### Functional annotation of inferred *L. stagnalis* genes

We first used TransDecoder v5.5.0 (Haas et al., 2013) to retrieve CDS and amino acid sequences for each assembled transcript from the *L. stagnalis* reference genome based on the merged annotation file generated by Stringtie. We applied two different methods to annotate inferred *L. stagnalis* genes and combined the annotated information. The first method was based on BLASTP searches against NCBI RefSeq amino acid sequences of nine species closely related to *L. stagnalis*. These species included: *Biomphalaria glabrata, Aplysia californica, Lottia gigantea, Pomacea canaliculata, Octopus bimaculoides, Octopus vulgaris, Crassostrea virginica, Crassostrea gigas*, and *Mizuhopecten yessoensis*. The second method was to search for the presence of Pfam domains in the inferred *L. stagnalis* amino acid sequences using the “hmmscan” tool in HMMER3 (Mistry, Finn, Eddy, Bateman, & Punta, 2013). The BLASTP and Pfam search results were integrated into the annotation of predicted *L. stagnalis* open reading frames (ORFs) using TransDecoder-v5.5.0 (Haas et al., 2013).

### Gene Ontology annotation of inferred genes in *L. stagnalis*

We used predicted protein sequences of *L. stagnalis* with at least 100 amino acids for Gene Ontology (GO) annotation using Blast2GO (Conesa et al., 2005). This annotation analysis was based on homology searches against the Mollusca phylum, *Caenorhabditis elegans, Drosophila melanogaster*, and *Homo sapiens* using the latest reference protein database (refseq_protein v5). We used an e-value threshold of 1.0E-3, top 20 blast hits, word size 6, and HSP length cut-off of 33. GO annotation was based on the latest GO version (2020.06). For GO enrichment analysis, Fisher’s exact test was used in combination with a False Discovery Rate (FDR) correction for multiple testing (FDR < 0.05). R (Team, 2017) package ggplot2 was used to plot results of GO enrichment analysis.

### Identification of previously cloned genes in *L. stagnalis*

We searched for previously cloned genes in *L. stagnalis* from the NCBI nucleotide database (as of January 2021) to evaluate the completeness of our transcriptome assembly. Only cloned genes that were supported by published studies were selected. The list of previously cloned genes in *L. stagnalis* (NCBI ID, gene names and references) is provided in **Supplemental Table 2**.

### Differential gene expression analysis and validation of gene expression by real-time qPCR

Differential expression (DE) analysis was carried out using DESeq2 (Love, Huber, & Anders, 2014) based on raw read counts retrieved by the featureCounts package of Subread v1.5.0 (Liao, Smyth, & Shi, 2014). The results of DE analysis by DESeq2 are shown in **Supplemental file 1**.

We validated the differential gene expression through real-time quantitative polymerase chain reactions (RT-qPCR). cDNA synthesis was performed from gDNA-removed RNA samples using SuperScript IV VILO Master Mix (Invitrogen, 11766050) following the manufacturer’s instructions. SYBR Green PCR Master Mix (Applied Biosystems, 4309155) was used for RT-qPCR in a QuantStudio 5 Real-Time PCR System (ThermoFisher). Primers are listed in **Supplemental Table 3**. Primer set efficiency values ranged from 95.85-104.26%, and R^2^ values were 0.9807-0.9999. Two negative controls were used: qPCR without reverse transcription and no template controls. Relative gene expression was normalized to reference gene *β-tubulin*. The final qPCR product was also sequenced to ensure the correct *innexin* paralog was amplified.

Because of the wide range of primer efficiencies, relative gene expression was calculated via the Common Base Method (Ganger, Dietz, & Ewing, 2017) and normalized to reference gene *β-tubulin*. Analysis of variance (ANOVA) was used to determine statistically significant differences in gene expression at p<0.05, and Tukey’s HSD *Post-hoc* test was used when appropriate.

## Results

### *L. stagnalis* CNS transcriptome sequencing, assembly and gene annotation

RNA-Seq was performed using CNS samples from young (3 months old), adult (6 months old), and old (18 months old) snails (**Figure 1A**), with four biological replicates in each group and ten snails in each replicate. Our RNA-Seq data provides a good sequencing depth, with a total number of reads ranging from 55,601,129 (56M) to 69,121,300 (69M). The average overall alignment rate is ∼74% (**Supplemental Table 4**). A total of 61,994 transcripts from 42,478 genes are identified. 37,661 of those genes encode for proteins of at least 100 amino acids. To provide functional annotations for inferred *L. stagnalis* genes, we retrieved proteomic sequences of nine molluscan species from the NCBI RefSeq database (See Materials and Methods). Our transcript assembly and gene function annotation provide the first genome annotation for *L. stagnalis* (provided as **Supplemental file 2** in gff3 format).

**Figure 1.**
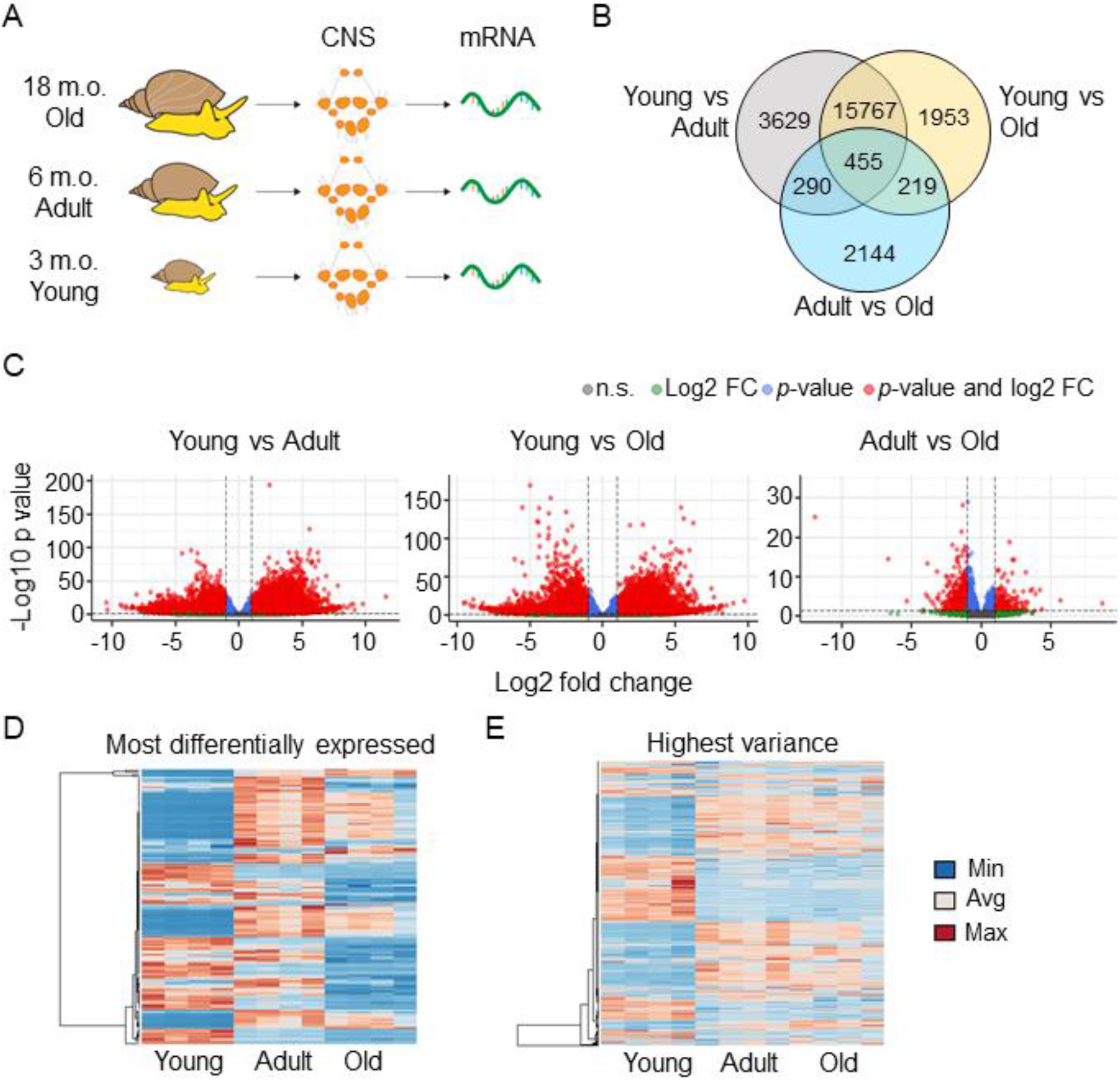
Pairwise comparisons of transcriptomes reveal a specific pattern of gene expression in the CNS of young *L. stagnalis*. A) For transcriptome analysis, mRNA was extracted from the CNS of young (3 months old), adult (6 months old), and old (18 months old) snails. For each age group, four different biological replicates (*n* = 4), each with mRNA samples from the CNS of 10 snails, were used. B) Venn diagram showing the significantly differentially expressed genes in each pairwise comparison and the overlap among them. The diagram clearly shows that young CNS transcriptome has more significantly differentially expressed genes compared to adult and old. (FDR adjusted *p*-value *p* < 0.05; log2 fold change > |1|). C) Volcano plot of each pairwise comparison. The plots are color-coded based on the log2 fold change (green dots), and corrected *p*-value (blue dots). Red dots highlight genes that are significant and whose expression is highly changed (FDR adjusted *p*-value *p* < 0.05; log2 fold change > |1|). Volcano plots of the comparison between adult and old CNS transcriptome shows less differentially expressed genes, by either *p*-value or log2 fold-change, compared to the pairwise comparison of young versus adult or old transcriptome. D, E) Heatmap of the most differentially expressed genes in all pairwise comparisons (FDR adjusted *p*-value *p* < 0.05) and the genes with the highest variance (top 2000 genes), respectively. Both heatmaps show a distinct pattern of gene expression in the CNS transcriptome of young snails compared to adult and old.

### Transcriptional clustering pattern in CNS of young, adult, and old *L. stagnalis*

Our principal component analysis (PCA) of the 12 transcriptomes (three age groups, four replication samples per age group) form three major clusters (**Supplemental Figure 1A**), corresponding to the three age groups of samples. The majority of biological replicates cluster together, suggesting that expression profiles are more similar in animals belonging to the same-age cohort. The first principal component (PC1), which accounts for 68.12% of the variance in the data, provides separation between young and the other two groups (adult and old). The second principal component (PC2) only accounts for 7% of the variance, serving as a discriminator between adult and old transcriptomes. These patterns suggest that there are constitutive differences in transcriptomes between young and adult/old CNS, while adult and old CNS transcriptomes are more similar to each other. These results are consistent with pairwise Pearson correlations between these samples (**Supplemental Figure 1B)**.

To identify genes that are differentially expressed (DE) during the development of the CNS, we conducted pairwise comparisons of transcriptomes for the three groups of samples: young vs. adult, young vs. old, and adult vs. old. We identify 20,141 significant DE genes between young and adult groups; 18,394 DE genes between young and old groups; and only 3,108 significant DE genes between adult and old groups (FDR adjusted *p*-value *p* < 0.05; log2 fold change > |1|) (**Figure 1B,C**). Interestingly, only 455 DE genes are present in all comparisons. Together, these analyses suggest that most changes in CNS development occur pronouncedly during transitions from young to adult, and less changes occur during transitions from adult to old.

### Analysis of the DE genes confirms distinct gene expression patterns in young, adult, and old *L. stagnalis*

Next, we selected the genes that are the most DE, based on their adjusted p-values in each pairwise comparison. We first sorted the top 100 DE genes and then further refined by selecting only the sequences with at least 100 codons; a total of 143 most DE genes are used. Heatmap analysis of FPKM expression shows a different expression pattern among age groups (**Figures 1D,E**). Specifically, consistent with the above principal component analysis, individual replicates exhibit very similar regulation patterns within the same age cohort (**Figure 1D**). Overall, the adult animal transcriptome shows more highly expressed genes, while around half of genes in young and the majority of genes in old exhibit low expression. The heatmap pattern also shows that 1) most genes with low expression in young animals are highly expressed in adult and remain high in old; 2) most highly expressed genes in young animals have low expression in adult and become further down-regulated in old animals; 3) only a few clusters contain genes whose expression increases from young to adult and then decreases from adult to old (**Figure 1D**). We also looked at the top 2,000 genes with the highest variance across all samples. The heatmap in **Figure 1E** shows 1) variance in DE gene expression, again, separates young transcriptome from transcriptomes of adult and old, and the highest variance occurs in the young animal transcriptome; 2) two big clusters of genes increase in expression from young to adult CNS transcriptome and retain relatively high expression in old; 3) two big clusters of genes decrease in expression from young to adult CNS transcriptome and further lower their expression in old; 3) a few small clusters of genes increase expression from young to adult and then decrease from adult to old. Together, these data indicate that there are distinct changes of transcript profiling across life stages of animals. Next, we sought to study what sets of DE genes and related pathways are involved in the expression patterns across different life stages.

### Gene ontology analysis

We performed GO annotation with Blast2Go (Conesa et al., 2005) for the 37,661 transcripts that encode proteins with at least 100 amino acids (see Materials and Methods). A total of 32,288 transcripts (or genes) were successfully annotated with GO terms. GO enrichment analysis was performed for DE genes in: all pairwise comparisons (455 genes), young compared to adult CNS transcriptome (20,141), young compared to old CNS transcriptome (18,394), and adult compared to old CNS transcriptome (3,108) (**Figure 2**). The enriched GO terms of DE genes in all pairwise comparisons include biological processes that are related to metabolism of nitrogen compounds, organic substances, macromolecules, and cellular macromolecules. As expected from the huge overlap among significant DE genes, GO enrichment from young compared to adult and young compared to old groups partially overlap. In these two comparisons, the common GO terms in biological process are related to gene expression (transcription by RNA polymerase II and positive or negative regulation of gene expression). GO terms in cellular components are related to mitochondrial and ribosome pathways (mitochondrial ribosome, mitochondrial matrix, mitochondrial membrane, cytosolic ribosome, and ribonucleoprotein complex). Finally, GO terms in molecular function are related to signalling receptor and signalling transduction pathways (signalling receptor activity, G protein-coupled receptor activity, and transmembrane signalling receptor activity) (**Figure 2**). The common enriched metabolic, mitochondrial, and ribosomal pathways suggest that transcripts engaged in cellular metabolic, energy production, and protein synthesis activity are actively regulated in the CNS of snails.

**Figure 2.**
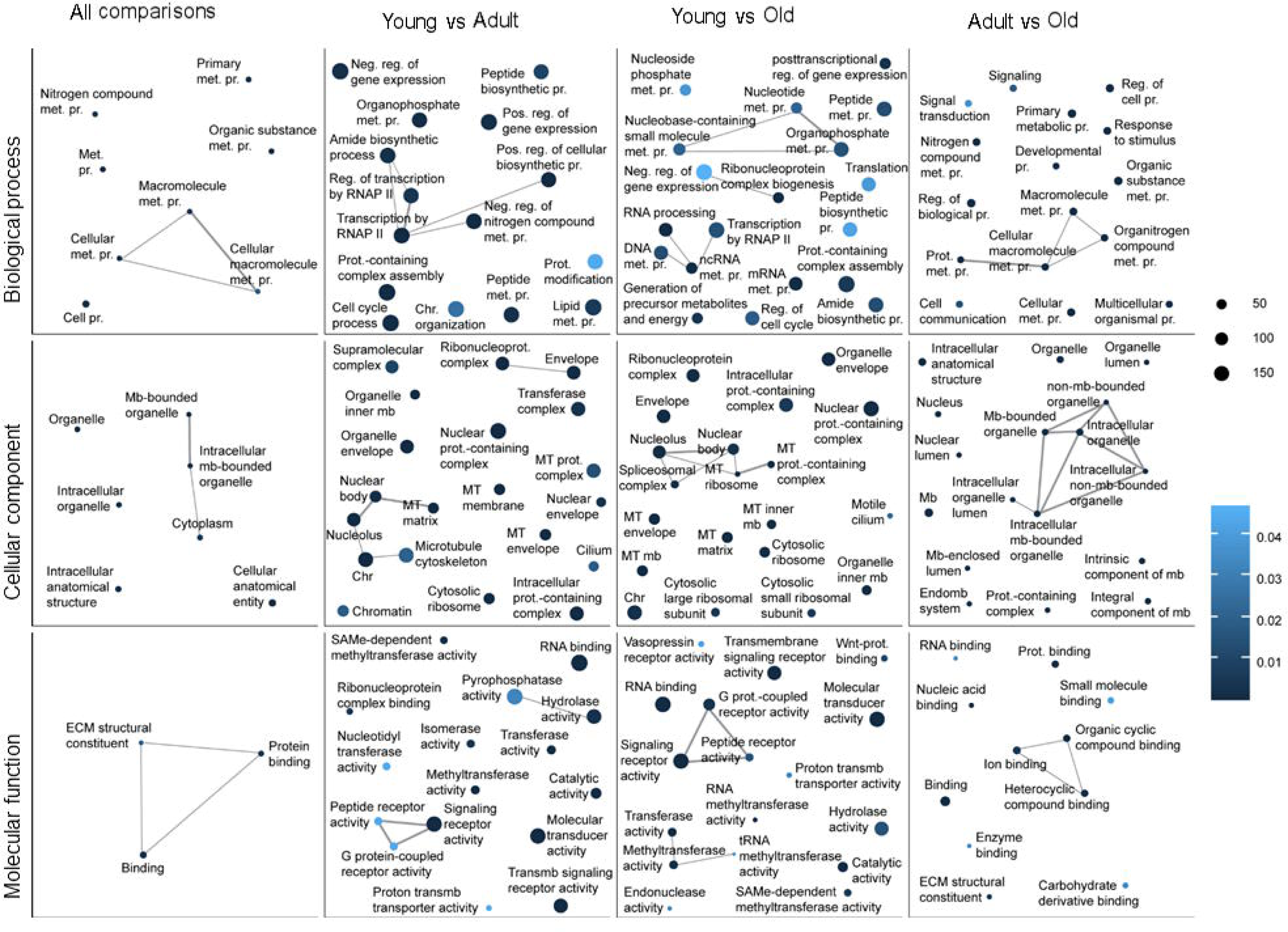
Gene Ontology (GO) enrichment analysis of differentially expressed genes. Representative enriched GO terms are shown as dots for each GO category (Biological process, Cellular component, or Molecular function). The significantly differentially expressed genes in all pairwise comparisons (All comparisons, 455 genes), in the comparison between young and adult CNS transcriptomes (Young vs. Adult, 20,141 genes), young and old CNS transcriptomes (Young vs. Old, 18,394 genes), or adult and old CNS transcriptomes (Adult vs Old, 3,108 genes) were used for the analysis. The dot size reflects the number of genes in the GO term for each significantly differentially expressed gene set; the dot colour represents the FDR-corrected *p*-value, with darker colours indicating lower *p*-values; the line size represents the degree of similarity. Abbreviations: met. = metabolic, pr. = process, neg. = negative, pos. = positive, reg. = regulation, mb = membrane, prot. = protein, chr = chromosome, RNAP II = RNA polymerase II, MT = mitochondrial, ECM = extracellular matrix.

To gain further insights into what specific sets of DE genes are changed from young to adult to old snail CNS, we analysed the FPKM expression of the top100 DE genes used in Figure 2D and examined their associated GO terms. A complete list of genes, their FPKM, descriptions, and GO terms are provided in **Supplemental table 5**. We found that genes significantly increased from young to adult and remain elevated in old animals, including those involved in receptor signalling activity (e.g. N-methyl-D-aspartate receptor NR1 and NR2, Mollusc insulin-related peptide MIP, and Notch 3 receptors), signalling transduction mechanism (serine/threonine protein kinase TAO1-like TAOK1, protein kinase A PKA, and serine/threonine-protein phosphatase 2A PIPA), synaptic vesicle proteins (e.g. synaptotagmin1 and 4), ion channels (e.g. voltage-gated K^+^ channels Kv2.1a KCNB1), metabolism (pyruvate kinase PKM-like isoform X3 PKM and phosphopractokinase-like PFKM), membrane/membrane bound organelles (e.g. reticulon-3-A like and fat cadherin), transcription and translation (poly [ADP-ribose] polymerase 14-like gene PARP and translation initiation factor eIF-2 EIF2B4), and peptides and peptide enzymes (Titin-like X2 TTN and Peptidase C1-like). Interestingly, there are only a few DE genes that are significantly increased from adult to old animals. These include monooxygenase/oxidoreductase active (CYP2U1), cysteine dioxygenase type (CD01), and endoglucanase E-4 like and A-like. Genes that are highly expressed in young and then significantly decrease in adult and old animals include ECM structure constituents (e.g. collagen 2A1, collagen 1A1, and fibril-forming collagen 2-chatin like). When comparing expression of genes in adult and old, we found that most genes exhibit significantly lower expression in old animals. These include stress and immune factors (dual oxidase 2-like DUOX2, oxidase activity cytochrome P450 CYP10, suppressor of cytokine signalling 2 SOCS2, and heat shock protein 60 HSP 60), Ca^2+^ binding (Ca^2+^/CaM-serine kinase CASK), protein ubiquitin (Myc binding protein MYCBP2 and ubiquitin-protein ligase E3A), and membrane and cellular entity (cadherin, adhesion G-protein coupled receptor L2 like GPCRL2, and disintegrin). Several above mentioned, stress-related genes and their expressions (FPKM) in young, adult, and old *L. stagnalis* CNS are shown in **Supplemental Figure 2**.

### qPCR studies confirm differential gene expression identified by RNA-Seq data

We next performed qPCR to validate gene expression revealed by RNA-Seq by taking advantage of our ongoing projects studying *L. stagnalis* gap junction *innexin* expression (Mersman et al., 2020). *Innexin* is the invertebrate analogue for the vertebrate *connexin*, both of which code for gap junction-forming proteins. Gap junctions are intercellular channels essential for direct and synchronized communication among cells (Ovsepian, 2017). Our transcriptome data detected three of the previously cloned *L. stagnalis innexin* genes: *Inx1, Inx4*, and *Inx5*. FPKM data showed that *Inx1* is the most abundant gene, followed by *Inx4*, and then *Inx5* in the CNS. The expression of these genes are upregulated in adult CNS and then maintain a comparable level in old CNS (**Figure 3A**). In order to compare the gene expression from our RNA-Seq data, we performed real-time qPCR from brain samples of young, adult, or old snails. As shown in **Figure 3A,B**, both RNA sequencing data and qPCR data show a similar pattern of expression over ages for each gene. More specifically, *Inx1* has a significantly lower expression level in young animals compared to adult and old (RNA-Seq FDR adjusted *p*-value young vs adult *p* = 5.95×10^−7^, young vs old *p* = 3.73×10^−6^; qPCR Turkey’s post-hoc test young vs adult *p* = 0.001 young vs old *p* = 0.009). *Inx4* also has similar trends in gene expression in both RNA-Seq and qPCR, but it is expressed significantly lower in the young animals only in the qPCR data, likely due to the increased sensitivity of the qPCR technique than RNA-Seq (RNA-Seq FDR adjusted *p*-value young vs adult *p* = 0.391, young vs old *p* = 0.395; qPCR Turkey’s post-hoc test young vs adult *p* = 0.003, young vs old *p* = 0.003). Finally, *Inx5* expression is also significantly lower in young snails compared to adult and old (RNA-Seq FDR adjusted *p*-value young vs adult *p* = 2.42×10^−11^, young vs old *p* = 0.000; qPCR Turkey’s post-hoc test young vs adult *p* = 0.001, young vs old *p* = 0.002). The concordance between RNA-Seq transcriptome expression and qPCR confirms the reliability of our RNA-Seq measurements.

**Figure 3.**
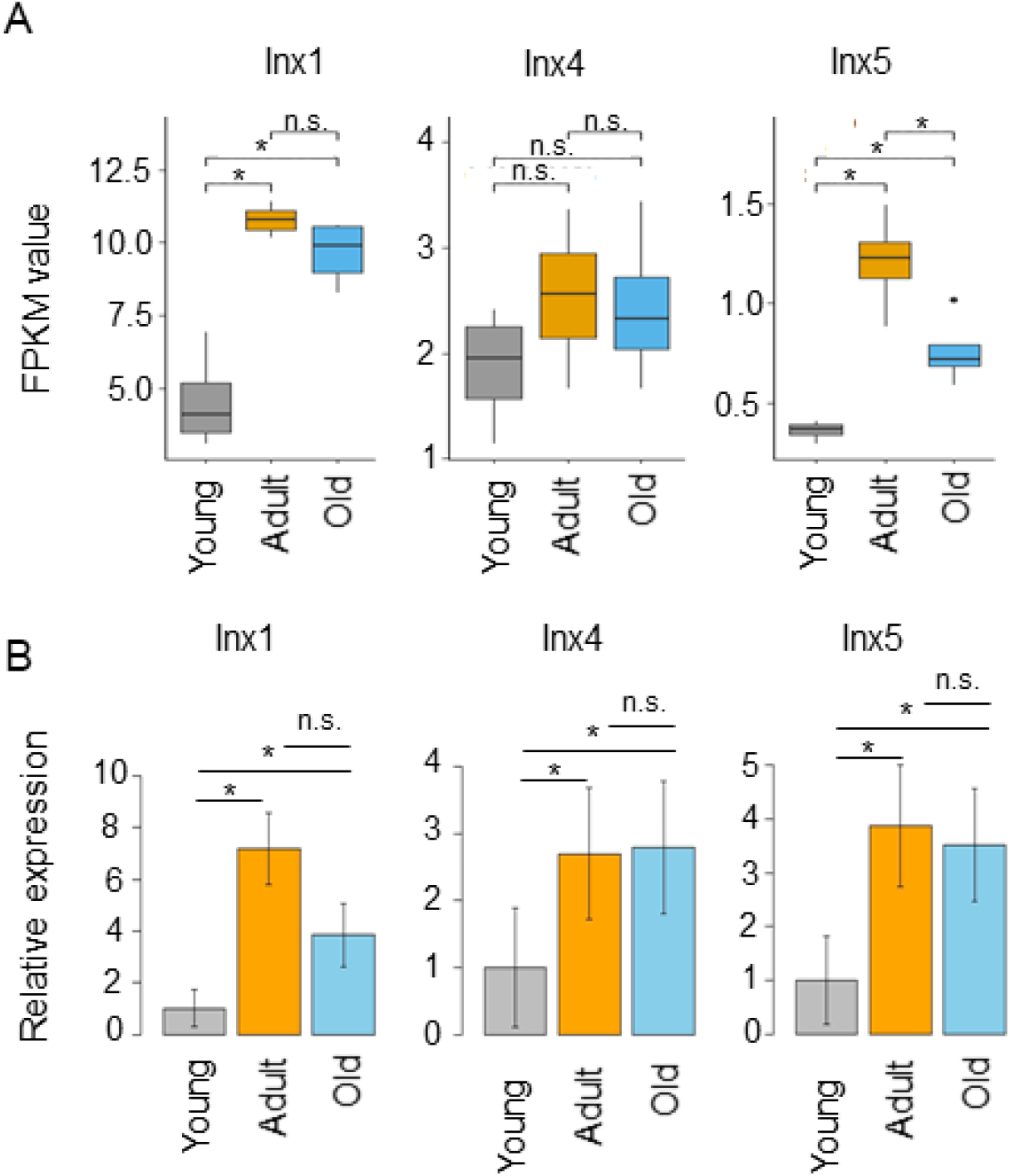
Confirmation of differential gene expression by real-time qPCR of *L. stagnalis Innexin* genes. A) RNA-Seq data reveals lower gene expression for innexins in young snails CNS compared to adult and old. B) real-time qPCR data show patterns of expression comparable to RNA-Seq data. These data show a general concordance of gene expression measured in our transcriptome data compared to qPCR data. * *p* < 0.05.

### *L. stagnalis* transcriptome assembly is further supported by the high coverage of other previously cloned genes

To further evaluate the quality and completeness of the transcriptome assembly, we tested the coverage of previously cloned genes from *L. stagnalis* in our transcriptome. According to the most recent NCBI nucleotide database, a total of 242 *L. stagnalis* genes have been previously cloned (**Supplemental Table 2**). There are 210 of these genes (87%) present in our transcriptome assembly, supporting that our transcriptome assembly from CNS samples cover most of the previously known protein-coding genes in *L. stagnalis*. Considering that only a portion of genes are expressed in brain tissues, it is expected that some previously cloned genes would not be detected by this RNA-Seq study.

We investigated the previously cloned genes among the aforementioned GO terms. We can find almost all of the previously cloned *L. stagnalis* acetylcholine receptor subunits (LnAChR) among the enriched signalling GO terms. Of the twelve previously cloned subunits, we found ten in our transcriptome. We further show that in the CNS transcriptome of adult and old snails, the LnAChR subunits H and F have the highest expression (van Nierop et al., 2006). Interestingly, in the CNS transcriptome of young snails, subunit G has the highest expression (**Figure 4A**). These data seem to suggest changes in LnAChR composition during CNS development. Moreover, two other synaptic receptors, the N-Methyl-D-aspartic acid receptor (NMDAR) and the serotonin receptor (5-HT receptor) have a significantly lower expression in the young compared to adult and old CNS transcriptomes (NMDAR: RNA-Seq FDR adjusted *p*-value young vs adult *p* = 6.35×10^−16^, young vs old *p* = 3.08E-17; 5-HT receptor: RNA-Seq FDR adjusted *p*-value young vs adult *p* = 9.90×10^−10^, young vs old *p* = 9.41×10^−5^) (**Figure 4B**). These data further suggest that, similar to humans, CNS synaptic development lasts after birth in snails, and a brain in a young individual is different from an adult or old brain, not only in ultrastructure, but also in phenotypes and transcriptome expression.

**Figure 4.**
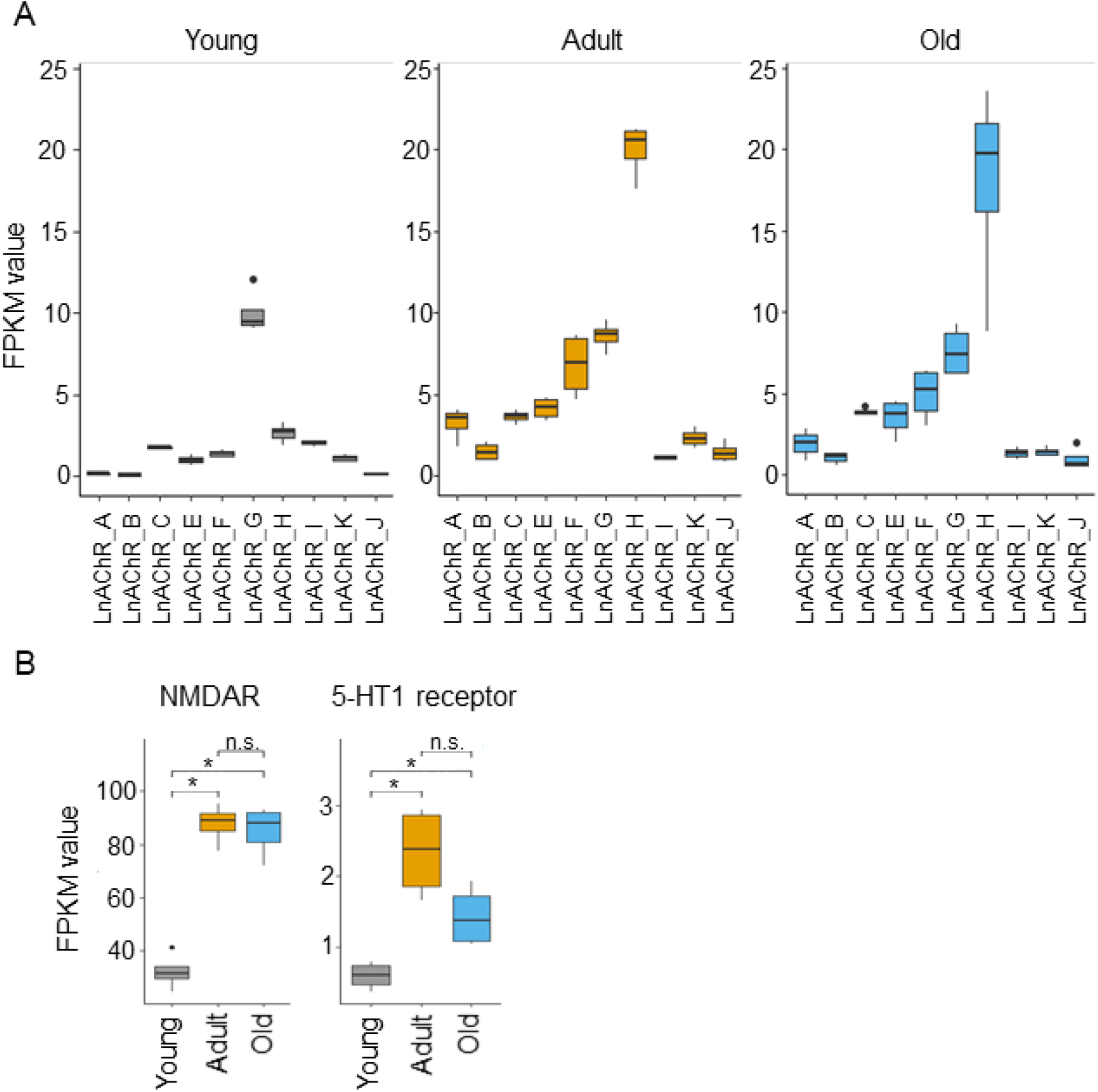
Differential expression of previously cloned synaptic receptor genes in *L. stagnalis*. A) FPKM expression of different *L. stagnalis* acetylcholine receptor subunits (LnAChR). Consistent with previous literature, the most highly expressed subunit in adult CNS transcriptome is the subunit H. Old snail CNS transcriptome has a similar pattern of LnAChR expression as adult snails. In young snails, though, the most highly expressed subunit is G. These data suggest that acetylcholine receptor subunits are specifically expressed at different life stages. B) Genes involved in synaptic transmission (NMDAR, N-Methyl-D-aspartic acid receptor; 5-HT receptor, and serotonin receptor) are significantly downregulated in young snails CNS compared to adult and old. The different patterns of expression for neurotransmitter receptors in young versus adult and old snails suggest that CNS synaptic development requires specific patterning to establish functional synapses throughout the span of *L. stagnalis* life. * *p* < 0.05.

Interestingly, of the known cloned genes matched in our transcriptome (210 genes), nine genes have significant differential gene expression in all pairwise comparisons (young vs adult, young vs old, adult vs old; **Figure 3A,B; Figure 5**). These include three genes coding for the molluscan insulin peptide (MIP, LSTA.21646), the enzyme tryptophan hydroxylase (LSTA.51), and the FMRFamide protein (LSTA.14894) that significantly increase in expression from young to adult to old snails (**Table 1, Figure 5**). Two genes coding for aquaporin channel protein (CoAQP1, LSTA.13986) and chitinase II (LSTA.17318) have an inverse pattern with significantly decreasing gene expression from young to adult to old animals (**Table 1, Figure 5**). Another three genes coding for a serum-dependent glutathione peroxidase (LSTA.693), *innexin5* (Inx5, LSTA.18425, as previously shown in **Figure 3A**), and another aquaporin channel protein (LsAQP1, LSTA.13985) are expressed higher in adult snails compared to young and old (**Table 1, Figure 5)**. Lastly, the gene coding for a voltage-dependent L-type calcium channel alpha-1 subunit was highly expressed in young snails compared to adult and old (**Table 1, Figure 5**).

**Table 1.** Significantly differentially expressed cloned genes.

| Gene | Locus ID | FPKM-Young | FPKM-Adult | FPKM-Old |
| --- | --- | --- | --- | --- |
| **MIP** | LSTA.21646 | 978.861 | 6055.743 | 7515.437 |
| **tryptophan hydroxylase** | LSTA.51 | 64.414 | 447.099 | 698.380 |
| **FMRFamide protein** | LSTA.14894 | 686.373 | 1042.714 | 1357.454 |
| **CoAQP1** | LSTA.13986 | 142.268 | 58.499 | 27.372 |
| **chitinase II** | LSTA.17318 | 5.811 | 5.348 | 3.259 |
| **serum-dependent glutathione peroxidase** | LSTA.693 | 2.669 | 31.388 | 18.185 |
| **Inx5** | LSTA.18425 | 0.361 | 1.208 | 0.761 |
| **LsAQP1** | LSTA.13985 | 115.796 | 176.112 | 69.177 |
| **voltage-dependent L-type calcium channel alpha-1 subunit** | LSTA.19085 | 1.164 | 0.757 | 0.759 |

**Figure 5.**
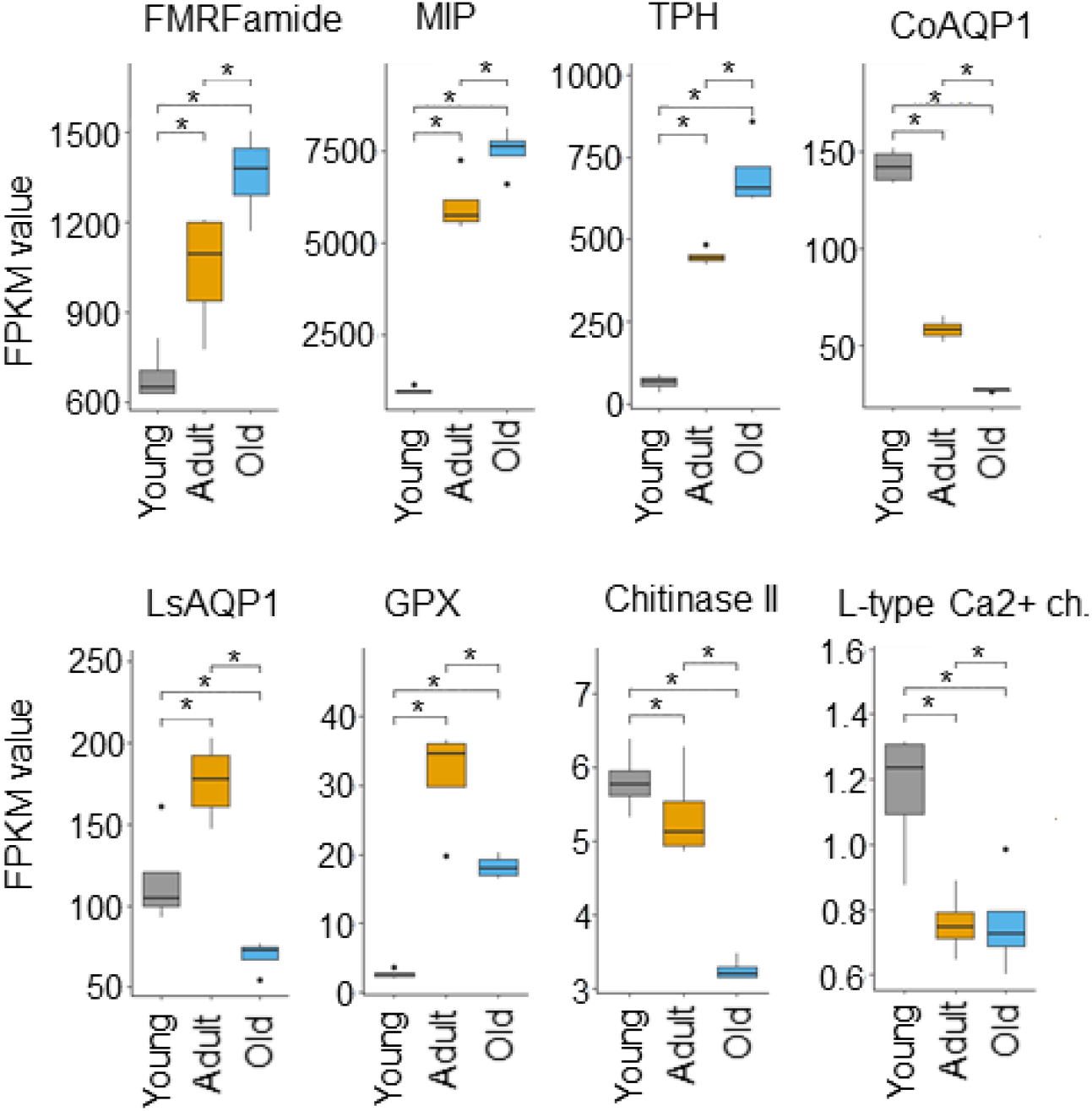
Cloned genes with significantly different expression at all pairwise comparisons. The figure shows eight of the nine genes (*Inx5* is shown in Figure 3A), among known cloned genes, that show significant changes in expression in all pairwise comparisons (young vs adult, young vs old, adult vs old) (FDR adjusted *p*-value *p* < 0.05). Differential expression of known genes suggests regulation of related pathways (e.g. long-term memory formation for MIP) in the *L. stagnalis* central nervous system at different ages. MIP, molluscan insulin peptide; TPH, tryptophane hydroxylase; AQP1, aquaporin1; GPX, glutathione peroxidase. * *p* < 0.05.

### Genes involved in diseases are upregulated in the CNS transcriptome of adult and old animals

Lastly, a recent paper using both *A. californica* and *L. stagnalis* has provided the cloned sequences for several genes involved in neurodegenerative disorders like Huntington disease, Parkinson’s disease (PD), and Alzheimer’s disease (AD) (Fodor, Urban, et al., 2020). Identification and expression analysis of these genes in the *L. stagnalis* transcriptome is promising for the use of *L. stagnalis* as a model for studying neurodegenerative diseases. In our transcriptome, we discovered that the expression of these genes change with age. More specifically, Parkinson’s disease protein 7 (*PARK7*), huntingtin, choline acetyltransferase, and presenilin genes are all upregulated in the CNS of adult and old compared to young animals (**Supplemental Figure 3**).

In addition to these well-recognized genes, there are several other genes that are upregulated in adult and old animals compared to young animals. These include Arginase 1 (ARG1, linked to many human diseases), reticulons (linked to AD and amyotrophic lateral sclerosis) (Yang & Strittmatter, 2007), proton myo-inositol cotransporter (SCL2A13, linked to AD) (Teranishi et al., 2015), Rab GDP dissociation inhibitor (linked to mental retardation) (Ishizaki et al., 2000), and several tumor genes such as tumor protein D52 (TPD 52), tumor suppressor gene e-cadherin like, and protein phosphatase2A (PP2A); a few of these genes and their expressions in young, adult, and old *L. stagnalis* are shown in **Supplemental Figure 3**). Expression of these disease-related genes in *L. stagnalis* provides a unique opportunity for using *L. stagnalis* as a model system to study these genes.

## Discussion

*L. stagnalis* has served as a unique model organism for the study of neural networks, neuronal development, and synapse formation (Getz et al., 2018; Haque et al., 2006; Swinton et al., 2019) due to its simple, well-characterized, and easily accessible CNS. In addition, it has recently emerged as a useful model for studying brain aging and neurodegenerative diseases (Fodor, Urban, et al., 2020; Hermann et al., 2020; Maasz et al., 2017). Here, we generated datasets that allow for the first in-depth look at transcriptome changes in gene expression of *L. stagnalis* CNS from young (3 month), adult (6 month), and old (18 month) animals. Our study identifies new *L. stagnalis* sequences, with a good read depth of up to 69 million total fragments (150 bp paired-end reads); moreover, we took advantage of the Blast2GO bioinformatics platform to provide gene annotation and gene ontology (GO) annotation for over 30,000 sequences. This study also reveals temporal dynamics of transcriptional profiling and key DE genes/pathways in *L. stagnalis* CNS at multiple time points of the animal’s life span. Such information will be instrumental for future age-related phenotypical analyses in single cells, neuronal networks, and whole animals.

Knowledge about age-associated transcript changes can improve our understanding of how intrinsic profiling plays a role in influencing anatomical, physiological, and pathophysiological properties in animals and human at different life stages. Transcriptomic analyses of human brains at different ages have shown that the majority of protein-coding genes are spatiotemporally regulated, and the transcriptional differences are most pronounced during early development (Kang et al., 2011). Similarly, our data reveal that constitutive differences in transcriptomes exist between young and adult CNS, while adult and old CNS exhibit fewer differences in transcripts. In the rat hippocampus, 229 genes were reported to be linearly up- or down-regulated across the lifespan of a healthy animal (Shavlakadze et al., 2019). Previous studies of transcriptomic profiling of sensory neurons from *A. californica* at 8 (matured), and 12 (aged) months reported that half of the genes were up- and half of the genes were down-regulated between the two cohorts of neurons (Greer et al., 2018). Our data in this study demonstrates that in *L. stagnalis* CNS, there are genes exhibiting linear up- or down-regulation from young to adult to old (Figure 1D), but there are also many genes that are regulated in a non-linear manner. For example, some genes are upregulated from young to adult and appear to either maintain a comparable level of expression or significantly decreased expression in old animals as shown in **Figures 3-5** and **Supplemental Figure 2-3**. The linear or non-linear expression of genes across animals at different ages and in different species may highly correlate with age- and species-specific functions of these genes. Interestingly, the principal component analysis revealed that the majority of *L. stagnalis* (**Supplemental Figure 1**) biological replicates clustered together. The same clustering pattern was found in *A. californica* (Greer et al., 2018) suggesting that individuals of these two species in the same age group share similar transcript profiles. These results suggest that age is an important, determinizing factor for transcriptional profiling between individuals. Together, these data support that whole transcriptome comparison can serve as a valuable tool for discovering age-specific genes, and these mollusc organisms could serve as useful models for studying age-related molecular basics of brain development and aging.

Our gene ontology analyses indicate that the majority of DE genes occur between young and adult CNS, and the DE genes are enriched in metabolic processes, gene expression, and mitochondrial, ribosome, and signalling receptor pathways. This finding is consistent with a recent study of adult *L. stagnalis* CNS transcriptome compared to several other adult organisms used in neurobiology including *Mus musculus* (mouse), *Danio rerio* (zebrafish), *Xenopus tropicalis, C. elegans*, and *D. melanogaster* (*Dong et al., 2021)*. Specifically, this study focused on the annotation of the top 20 expressed transcripts in these models. The authors revealed an abundance of transcripts involved in energy production, protein synthesis, and signalling transduction of adult CNS, indicating evolutionally conserved roles of these pathways in mature adult animals of both invertebrates and vertebrates. Because of the importance of these pathways in adult animals, it is not surprising that previous studies have primarily focused on cloning genes involved in these pathways. Indeed, among many cloned genes in our *L. stagnalis* transcriptome, we can detect DE genes encoding proteins involved in these pathways. For example, our RNA-Seq data identified ten of the twelve *L. stagnalis* LnAChR subunits that have been previously cloned and sequenced (van Nierop et al., 2006). Interestingly, we provide evidence that these subunits are differentially expressed in the snail’s CNS at different ages: the subunit H is most highly expressed in adult and old, followed by the F subunit, while the G subunit is most highly expressed in young snails. The expression of LnAChR subunits in adult snails is consistent with literature showing that subunits H and F together account for approximately half of LnAChR expression in the *L. stagnalis* CNS as shown by *in situ* hybridization (ISH) (van Nierop et al., 2006). Studies in rat brain have also shown that various nAChR subunits are expressed at different ages and in different brain areas (Cimino et al., 1995; X. Zhang, Liu, Miao, Gong, & Nordberg, 1998). Studies comparing primates and humans to rodent brains have shown that the expression of nAChR subunits is conserved in some brain areas but not others (Zoli, Pistillo, & Gotti, 2015). Furthermore, molluscan and other invertebrate species are known to have not only excitatory, sodium-selective nAChRs, but also inhibitory, chloride-selective nAChRs (van Nierop et al., 2005). Considering the properties of cation or anion conductance of LnAChR subunits, we can appreciate the importance of differential expression of these subunits across the lifespan of *L. stagnalis* for maintaining excitability homeostasis. Specific pharmacological properties have been demonstrated for nAChRs composed of different subunits in both mammalian (Papke, Dwoskin, & Crooks, 2007; Zoli et al., 2015), and invertebrate models (Lansdell, Collins, Goodchild, & Millar, 2012). Our transcriptome data suggest that the expression pattern or properties of LnAChRs might be important in CNS development and function. Therefore, it would be interesting to investigate the pharmacological properties of the different LnAChR subunits and their spatiotemporal expression and function in the CNS of *L. stagnalis* in the future.

Among other previously cloned neurotransmitter receptors, we found that the N-Methyl-D-aspartic acid (NMDA) and the serotonin (5-HT) receptors are differentially expressed when comparing the CNS of young and adult/old snails (**Figure 4B**). Studies in both human (Bar-Shira, Maor, & Chechik, 2015; Law et al., 2003) and rat (Monyer, Burnashev, Laurie, Sakmann, & Seeburg, 1994) have shown differential expression of the NMDAR subunits NR1 and NR2 at different ages. Similarly, 5-HT receptors have been shown to be differentially expressed during human early postnatal development and into adulthood (Bar-Shira et al., 2015; Lambe, Fillman, Webster, & Shannon Weickert, 2011). Importantly, aberrant expression and/or function of NMDA and 5-HT receptors have been associated with neurodevelopmental disorders such as schizophrenia and autism (Carlsson, 1998; du Bois & Huang, 2007; Ju & Cui, 2016; Seshadri, Klaus, Winkowski, Kanold, & Plenz, 2018; Sodhi & Sanders-Bush, 2004; Xia et al., 2018). These changes in neurotransmitter receptor expression at different ages suggest ongoing synaptic development or synaptic diversification when *L. stagnalis* CNS progresses from young to a fully matured stage. The significant up-regulation of genes encoding these transmitter receptors as well as synaptotagmin, gap junctions, ion channels, FMRFamide and Mollusc insulin-related peptides (**Supplemental Table 5** and results) clearly indicates the active engagement of intercellular communication and synaptic plasticity in these adult animals. Transcript regulation of synaptic genes may reflect animal behavioural changes; compared to young, adult and old animals normally exhibit more vigorous and diverse behaviours including reproduction, feeding, locomotion, respiration, and associative learning, for which the above-mentioned synaptic machinery components play major roles (Dong et al., 2021; Ha, Kohn, Bobkova, & Moroz, 2006; Hoek et al., 2005; Ito et al., 2012; Kojima et al., 2015; Mersman et al., 2020; Murakami et al., 2013; Yeoman et al., 1994).

Neural communication relies on both the transmitter receptor-mediated chemical synapse and the gap junction-mediated electrical synapse (Pereda, 2014); however, the latter is severely understudied in *L. stagnalis* due to the lack of genetic information for gap junction forming innexins. Recently, eight *innexin* genes, *Lst Inx1-Lst Inx 8*, have been sequenced and cloned in *L. stagnalis* by our lab (Mersman et al., 2020). We mapped three of these genes, *Lst Inx1, Lst Inx4*, and *Lst Inx5*, in our transcriptome data. The other *innexin* genes might have expression levels that are too low to be measured in our transcriptome data, or the genome assembly we used for gene mapping might have distributed those sequences on different scaffolds. If we look at the three *innexin* genes that we were able to detect, both transcriptome and RT-qPCR confirm differential expression at different ages (**Figure 3A-B**). Studies in vertebrates have shown that specific connexin composition determines gap junction channel properties (Beyer, Lipkind, Kyle, & Berthoud, 2012; Rackauskas, Neverauskas, & Skeberdis, 2010). Moreover, a study in the invertebrate *D. melanogaster* showed that two innexins, shakB and ogre, are not interchangeable, as they fail to rescue the other’s mutant (Curtin, Zhang, & Wyman, 2002). The differential expression of *innexin* genes in *L. stagnalis* further suggests that gap junction channels formed by different innexin proteins might have a specific role and are, hence, differentially expressed at different ages.

In addition to previously known genes, it is interesting to note a few unstudied genes in the *L. stagnalis* transcriptome that exhibit age-specific expression patterns across life stages (as described in results). For example, genes related to oxidative stress and immunity response are either up- or down-regulated in old animals when compared to young and adult animals (**Supplemental Figure 2**). These include cytochrome P450 (CYP2U1 and CYP10), dual oxidase 2 (DUOX2), and suppressor of cytokine signalling 2 (SOCS2). The cytochromes P450 (CYPs) constitute a large superfamily of hemeproteins that are largely involved in the oxidative metabolism of environmental (xenobiotics such as drugs and pesticides) or endogenous (steroid hormones, fatty acid, etc) compounds (Dhers, Ducassou, Boucher, & Mansuy, 2017; Montellano, 2015). Both CYP2U1 transcripts and proteins are widely expressed in various brain regions of human and rats and are involved in the metabolism of fatty acids and xenobiotics in the brain (Dhers et al., 2017). Interestingly, CYP10 has been cloned in *L. stagnalis* and found to be abundantly expressed in the female gonadotropic hormone producing dorsal bodies (Teunissen, Geraerts, van Heerikhuizen, Planta, & Joosse, 1992). DUOX are oxidoreductase enzymes that catalyse the synthesis of reactive oxygen species (ROS) molecules including the anion superoxide (O_2_^-^) and hydrogen peroxide (H_2_O_2_). DUOX as well as the previously cloned GPX (**Figure 5**) have recently been demonstrated to be DE in *L. stagnalis* transcriptome in response to ecoimmunological challenges (Seppala et al., 2021). SOC2 acts as a negative feedback inhibitor for a variety of cytokine signalling in both vertebrates and invertebrates (Wang, Wangkahart, Secombes, & Wang, 2019; Y. Zhang et al., 2010). Because of the significant regulation of these oxidative stress and immune defense genes across life stages, it is critical to study their roles in animal health, aging, and diseases in future studies.

Finally, our transcriptomic data also revealed changes in disease-relevant genes associated with neurodegeneration, aging, and cancer, thus affording a unique opportunity to study cellular and molecular functions of these genes by taking advantage of the simplicity of *L. stagnalis* CNS. In addition, sequenced homologs of several genes known to be involved in aging and neurodegenerative disease (e.g. Parkinson’s disease protein 7 (*PARK7*), huntingtin, presenilin1, and choline acetyltransferase (AChAT) (Fodor, Urban, et al., 2020), among others) have recently become available. Our data reveals that in *L. stagnalis*, the expression of these genes is upregulated in adult and old CNS compared to young CNS. In addition to these genes, in the present study, we have discovered several other disease-related genes (Supplemental Table 5 and Supplemental Figure 3). Firstly, the membrane proton myo-inositol cotransport (SLC2A13) increases expression in the CNS of adult and old compared to young *L. stagnalis*. Similar to presenilin1, proton myo-inositol cotransport is found to be a novel gamma-secretase associated protein that selectively regulates Aβ production (Teranishi et al., 2015). Secondly, the present study reveals increased expression of reticulons which have been linked to AD and amyotrophic lateral sclerosis (ALS) (Yang & Strittmatter, 2007). Reticulons is a group of evolutionarily conserved proteins residing predominantly in the endoplasmic reticulum that promote membrane curvature, vesicle formation, and nuclear pore complex formation. Since all these proteins are potentially related to AD, it would be interesting to investigate their roles in learning and memory or aging in future studies. While mutation, deletion, or decrease in expression of these disease-related genes are the primary cause of diseases (Bird, Stranahan, Sumi, & Raskind, 1983; Domingo & Klein, 2018; Nance, 2017; Nikolac Perkovic & Pivac, 2019), the purpose for maintaining a high expression of these gene transcripts in the CNS of adult and old animals is not known. However, our results may partially indicate that the abundant expression of these disease-related genes could be a result of normal physiological requirements or the natural aging process of mollusc CNS.

## Conclusions

Overall, our RNA-seq study provided a much-needed *L. stagnalis* transcriptome assembly, with gene and GO annotation for more than 30,000 predicted genes. Furthermore, the analysis of CNS from different ages demonstrates the importance of this model for uncovering molecular insights in young, adult, and old life stages. This dataset will be useful for future discoveries of genes, expression profiling, and signalling pathways in different ages of animals. It also serves as a helpful resource for future annotation of genes and the genome of *L. stagnalis*.

## Acknowledgement

This work was supported by the National Science Foundation (1916563) and the Saint Louis University Start-up Fund awarded to Dr. Xu.

## Competing interests

The authors declare no competing and financial interests.

---

**Supplemental Figure 1.**
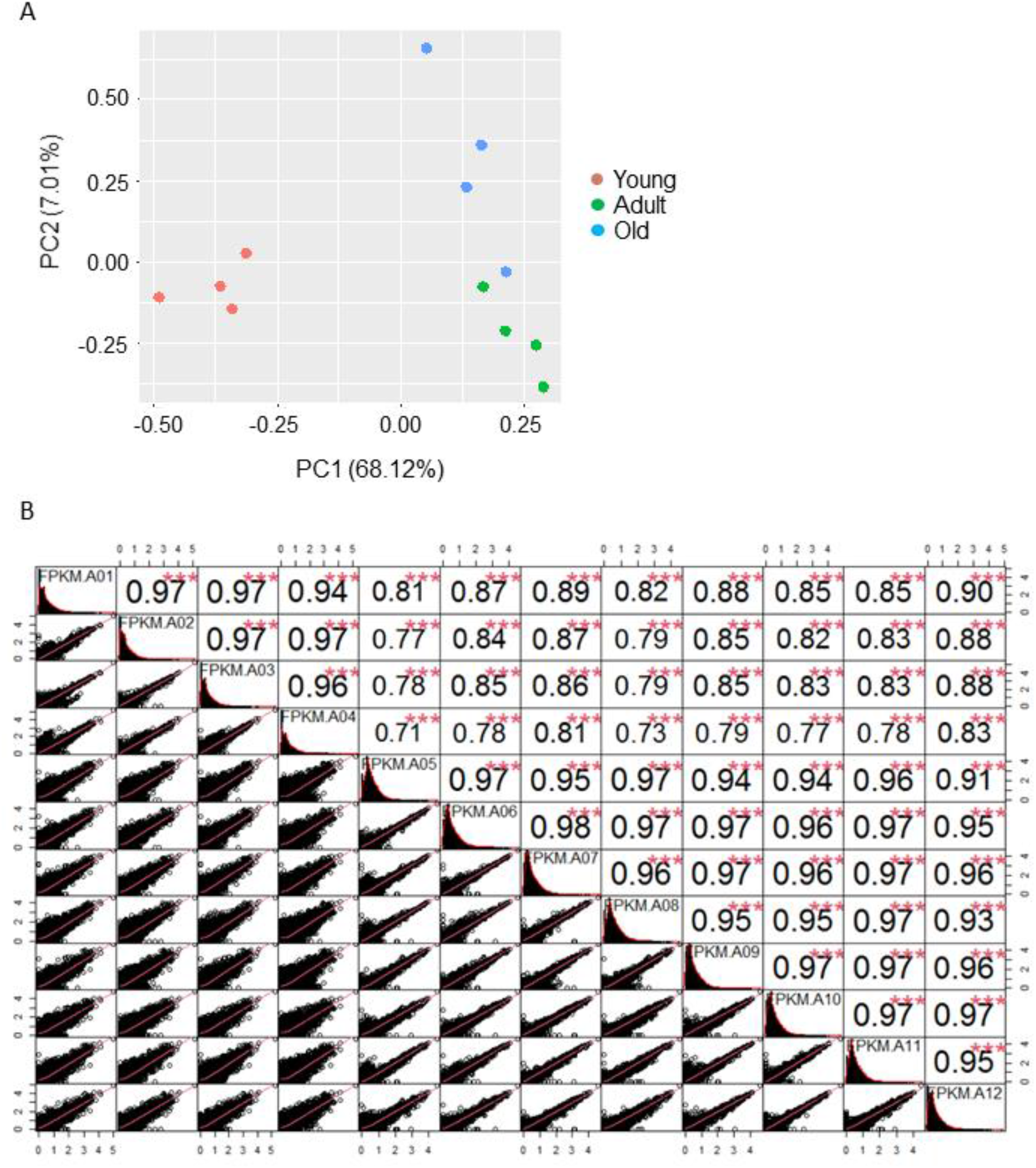
Data exploration of CNS transcriptomes reveal age-dependent separation of samples. A) Principal component analysis (PCA) shows a clear separation of different developmental age samples. The first component (x-axis) explains ∼68% of sample separation and clearly separates the CNS transcriptome of young snails from adult and old. The second component (y-axis) separates adult and old samples. B) Correlation analysis of gene expression profile shows high correlations among samples that belong to the same group (young snails CNS transcriptome, samples A01-A04; adult snails CNS transcriptome, samples A05-A08; old snails CNS transcriptome, samples A09-A12)

**Supplemental Figure 2.**
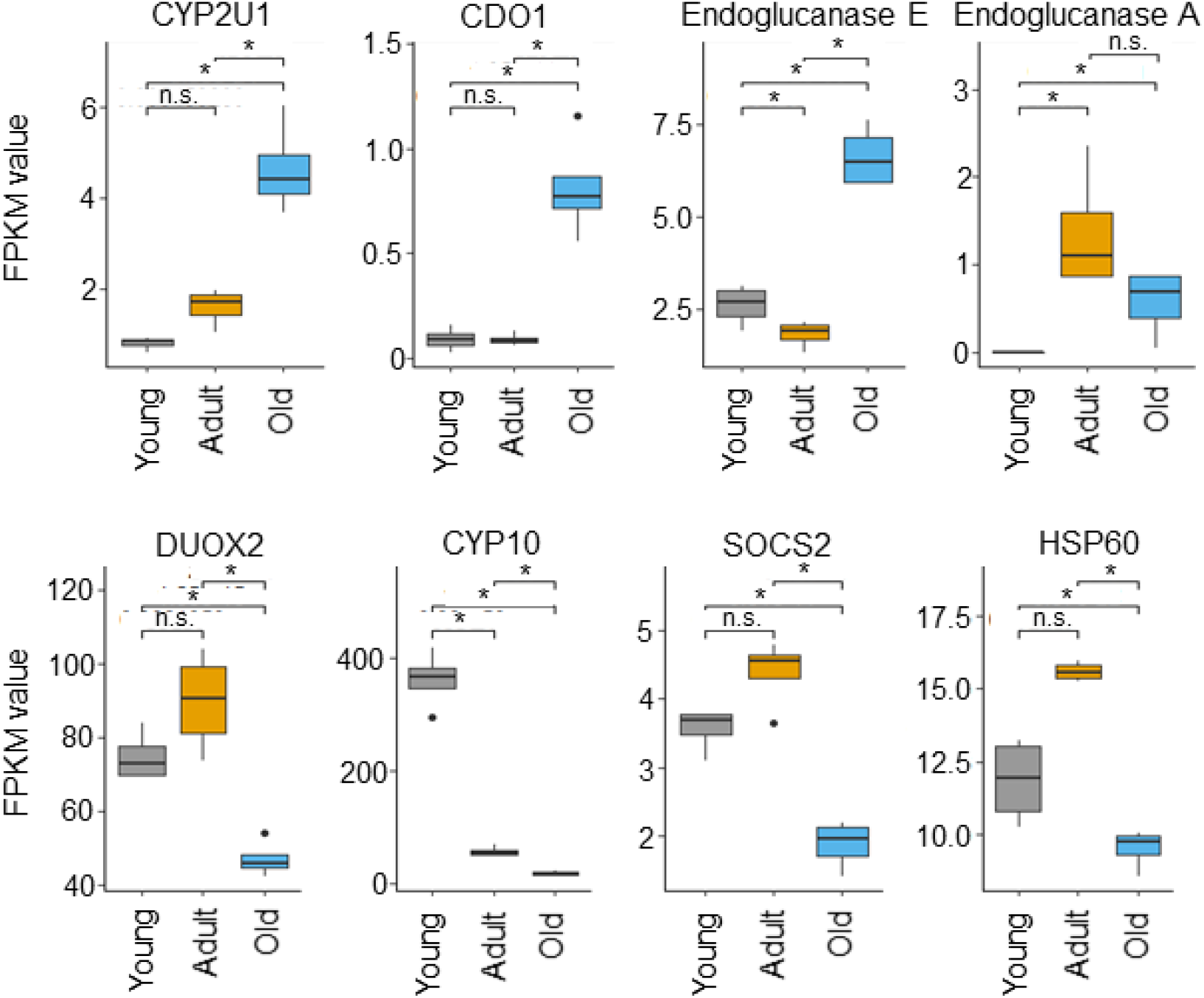
Differential expression of stress-related genes. Selected DE genes involved in stress and immune response demonstrate differential expression patterns. Cytochrome P450 (CYP2U1), cysteine dioxygenase type 1 (CDO1), Endoglucanase E, Endoglucanase A, dual oxidase 2 (DUOX2), cytochrome P450 (CYP10), suppressor of cytokine signaling 2 (SOCS2), and heat shock protein 60 (HSP60).

**Supplemental Figure 3.**
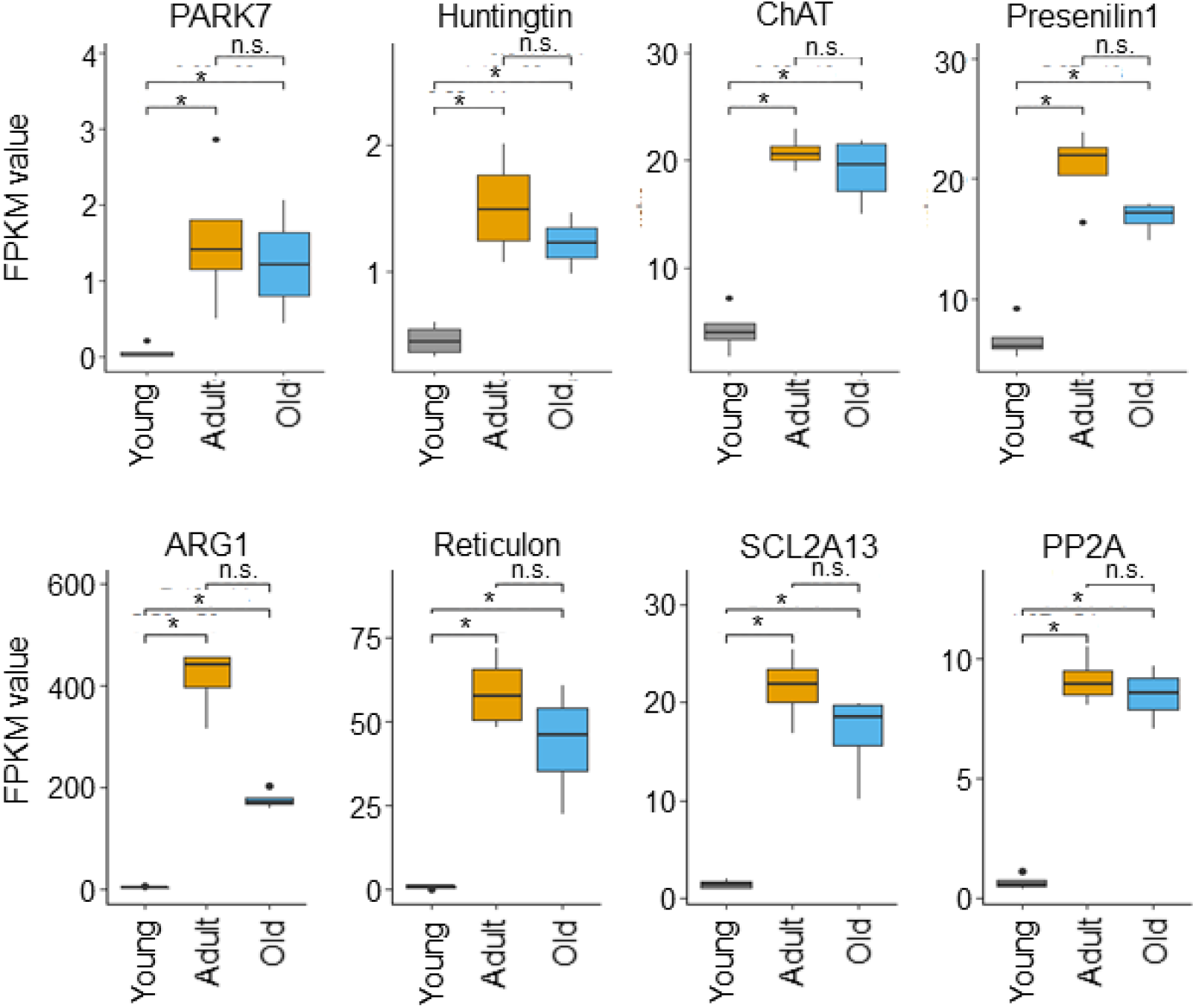
Differential expression of disease-related genes. Disease related genes have lowest levels of expression in young snails. Parkinson’s disease protein 7 (PARK7), huntingtin, choline acetyltransferase (ChAT), presenilin 1, arginase-1 (ARG-1), reticulon, membrane proton myo-inositol cotransporter (SCL2A13), and protein phosphatase 2A (PP2A).

## Notes

https://www.ncbi.nlm.nih.gov/bioproject/?term=PRJNA698985

### Competing Interest Statement

The authors have declared no competing interest.

## References

Abe, M., & Kuroda, R. (2019). The development of CRISPR for a mollusc establishes the formin Lsdia1 as the long-sought gene for snail dextral/sinistral coiling. Development, 146(9). doi:10.1242/dev.175976

Arundell, M., Patel, B. A., Straub, V., Allen, M. C., Janse, C., O’Hare, D., … Yeoman, M. S. (2006). Effects of age on feeding behavior and chemosensory processing in the pond snail, Lymnaea stagnalis. Neurobiol Aging, 27(12), 1880–1891. doi:10.1016/j.neurobiolaging.2005.09.040

Bar-Shira, O., Maor, R., & Chechik, G. (2015). Gene Expression Switching of Receptor Subunits in Human Brain Development. PLoS Comput Biol, 11(12), e1004559. doi:10.1371/journal.pcbi.1004559

Beyer, E. C., Lipkind, G. M., Kyle, J. W., & Berthoud, V. M. (2012). Structural organization of intercellular channels II. Amino terminal domain of the connexins: sequence, functional roles, and structure. Biochim Biophys Acta, 1818(8), 1823–1830. doi:10.1016/j.bbamem.2011.10.011

Bird, T. D., Stranahan, S., Sumi, S. M., & Raskind, M. (1983). Alzheimer’s disease: choline acetyltransferase activity in brain tissue from clinical and pathological subgroups. Ann Neurol, 14(3), 284–293. doi:10.1002/ana.410140306

Boeck, M. E., Huynh, C., Gevirtzman, L., Thompson, O. A., Wang, G., Kasper, D. M., … Waterston, R. H. (2016). The time-resolved transcriptome of C. elegans. Genome Res, 26(10), 1441–1450. doi:10.1101/gr.202663.115

Bouetard, A., Noirot, C., Besnard, A. L., Bouchez, O., Choisne, D., Robe, E., … Coutellec, M. A. (2012). Pyrosequencing-based transcriptomic resources in the pond snail Lymnaea stagnalis, with a focus on genes involved in molecular response to diquat-induced stress. Ecotoxicology, 21(8), 2222–2234. doi:10.1007/s10646-012-0977-1

Carlsson, M. L. (1998). Hypothesis: is infantile autism a hypoglutamatergic disorder? Relevance of glutamate - serotonin interactions for pharmacotherapy. J Neural Transm (Vienna), 105(4-5), 525–535. doi:10.1007/s007020050076

Chou, S. J., Wang, C., Sintupisut, N., Niou, Z. X., Lin, C. H., Li, K. C., & Yeang, C. H. (2016). Analysis of spatial-temporal gene expression patterns reveals dynamics and regionalization in developing mouse brain. Sci Rep, 6, 19274. doi:10.1038/srep19274

Cimino, M., Marini, P., Colombo, S., Andena, M., Cattabeni, F., Fornasari, D., & Clementi, F. (1995). Expression of neuronal acetylcholine nicotinic receptor alpha 4 and beta 2 subunits during postnatal development of the rat brain. J Neural Transm Gen Sect, 100(2), 77–92. doi:10.1007/BF01271531

Conesa, A., Gotz, S., Garcia-Gomez, J. M., Terol, J., Talon, M., & Robles, M. (2005). Blast2GO: a universal tool for annotation, visualization and analysis in functional genomics research. Bioinformatics, 21(18), 3674–3676. doi:10.1093/bioinformatics/bti610

Curtin, K. D., Zhang, Z., & Wyman, R. J. (2002). Gap junction proteins are not interchangeable in development of neural function in the Drosophila visual system. J Cell Sci, 115(Pt 17), 3379–3388.

Davison, A., & Blaxter, M. L. (2005). An expressed sequence tag survey of gene expression in the pond snail Lymnaea stagnalis, an intermediate vector of trematodes [corrected]. Parasitology, 130(Pt 5), 539–552. doi:10.1017/s0031182004006791

de Weerd, L., Hermann, P. M., & Wildering, W. C. (2017). Linking the ‘why’ and ‘how’ of ageing: evidence for somatotropic control of long-term memory function in the pond snail Lymnaea stagnalis. J Exp Biol, 220(Pt 22), 4088–4094. doi:10.1242/jeb.167395

Dhers, L., Ducassou, L., Boucher, J. L., & Mansuy, D. (2017). Cytochrome P450 2U1, a very peculiar member of the human P450s family. Cell Mol Life Sci, 74(10), 1859–1869. doi:10.1007/s00018-016-2443-3

Dodd, S., Rothwell, C. M., & Lukowiak, K. (2018). Strain-specific effects of crowding on long-term memory formation in Lymnaea. Comp Biochem Physiol A Mol Integr Physiol, 222, 43–51. doi:10.1016/j.cbpa.2018.04.010

Domingo, A., & Klein, C. (2018). Genetics of Parkinson disease. Handb Clin Neurol, 147, 211–227. doi:10.1016/B978-0-444-63233-3.00014-2

Dong, N., Bandura, J., Zhang, Z., Wang, Y., Labadie, K., Noel, B., … Feng, Z. P. (2021). Ion channel profiling of the Lymnaea stagnalis ganglia via transcriptome analysis. BMC Genomics, 22(1), 18. doi:10.1186/s12864-020-07287-2

du Bois, T. M., & Huang, X. F. (2007). Early brain development disruption from NMDA receptor hypofunction: relevance to schizophrenia. Brain Res Rev, 53(2), 260–270. doi:10.1016/j.brainresrev.2006.09.001

Feng, Z. P., Zhang, Z., van Kesteren, R. E., Straub, V. A., van Nierop, P., Jin, K., … Smit, A. B. (2009). Transcriptome analysis of the central nervous system of the mollusc Lymnaea stagnalis. BMC Genomics, 10, 451. doi:10.1186/1471-2164-10-451

Fodor, I., Hussein, A. A., Benjamin, P. R., Koene, J. M., & Pirger, Z. (2020). The unlimited potential of the great pond snail, Lymnaea stagnalis. Elife, 9. doi:10.7554/eLife.56962

Fodor, I., Urban, P., Kemenes, G., Koene, J. M., & Pirger, Z. (2020). Aging and disease-relevant gene products in the neuronal transcriptome of the great pond snail (Lymnaea stagnalis): a potential model of aging, age-related memory loss, and neurodegenerative diseases. Invert Neurosci, 20(3), 9. doi:10.1007/s10158-020-00242-6

Ford, L., Crossley, M., Vadukul, D. M., Kemenes, G., & Serpell, L. C. (2017). Structure-dependent effects of amyloid-beta on long-term memory in Lymnaea stagnalis. FEBS Lett, 591(9), 1236–1246. doi:10.1002/1873-3468.12633

Frias-Soler, R. C., Pildain, L. V., Parau, L. G., Wink, M., & Bairlein, F. (2020). Transcriptome signatures in the brain of a migratory songbird. Comp Biochem Physiol Part D Genomics Proteomics, 34, 100681. doi:10.1016/j.cbd.2020.100681

Ganger, M. T., Dietz, G. D., & Ewing, S. J. (2017). A common base method for analysis of qPCR data and the application of simple blocking in qPCR experiments. BMC Bioinformatics, 18(1), 534. doi:10.1186/s12859-017-1949-5

Getz, A. M., Wijdenes, P., Riaz, S., & Syed, N. I. (2018). Uncovering the Cellular and Molecular Mechanisms of Synapse Formation and Functional Specificity Using Central Neurons of Lymnaea stagnalis. ACS Chem Neurosci, 9(8), 1928–1938. doi:10.1021/acschemneuro.7b00448

Graveley, B. R., Brooks, A. N., Carlson, J. W., Duff, M. O., Landolin, J. M., Yang, L., … Celniker, S. E. (2011). The developmental transcriptome of Drosophila melanogaster. Nature, 471(7339), 473–479. doi:10.1038/nature09715

Greer, J. B., Schmale, M. C., & Fieber, L. A. (2018). Whole-transcriptome changes in gene expression accompany aging of sensory neurons in Aplysia californica. BMC Genomics, 19(1), 529. doi:10.1186/s12864-018-4909-1

Ha, T. J., Kohn, A. B., Bobkova, Y. V., & Moroz, L. L. (2006). Molecular characterization of NMDA-like receptors in Aplysia and Lymnaea: relevance to memory mechanisms. Biol Bull, 210(3), 255–270. doi:10.2307/4134562

Haas, B. J., Papanicolaou, A., Yassour, M., Grabherr, M., Blood, P. D., Bowden, J., … Regev, A. (2013). De novo transcript sequence reconstruction from RNA-seq using the Trinity platform for reference generation and analysis. Nat Protoc, 8(8), 1494–1512. doi:10.1038/nprot.2013.084

Haque, Z., Lee, T. K., Inoue, T., Luk, C., Hasan, S. U., Lukowiak, K., & Syed, N. I. (2006). An identified central pattern-generating neuron co-ordinates sensory-motor components of respiratory behavior in Lymnaea. Eur J Neurosci, 23(1), 94–104. doi:10.1111/j.1460-9568.2005.04543.x

Hermann, P. M., de Lange, R. P., Pieneman, A. W., ter Maat, A., & Jansen, R. F. (1997). Role of neuropeptides encoded on CDCH-1 gene in the organization of egg-laying behavior in the pond snail, Lymnaea stagnalis. J Neurophysiol, 78(6), 2859–2869. doi:10.1152/jn.1997.78.6.2859

Hermann, P. M., Lee, A., Hulliger, S., Minvielle, M., Ma, B., & Wildering, W. C. (2007). Impairment of long-term associative memory in aging snails (Lymnaea stagnalis). Behav Neurosci, 121(6), 1400–1414. doi:10.1037/0735-7044.121.6.1400

Hermann, P. M., Perry, A. C., Hamad, I., & Wildering, W. C. (2020). Physiological and pharmacological characterization of a molluscan neuronal efflux transporter; evidence for age-related transporter impairment. J Exp Biol, 223(Pt 2). doi:10.1242/jeb.213785

Heyland, A., Vue, Z., Voolstra, C. R., Medina, M., & Moroz, L. L. (2011). Developmental transcriptome of Aplysia californica. J Exp Zool B Mol Dev Evol, 316B(2), 113–134. doi:10.1002/jez.b.21383

Hoek, R. M., Li, K. W., van Minnen, J., Lodder, J. C., de Jong-Brink, M., Smit, A. B., & van Kesteren, R. E. (2005). LFRFamides: a novel family of parasitation-induced -RFamide neuropeptides that inhibit the activity of neuroendocrine cells in Lymnaea stagnalis. J Neurochem, 92(5), 1073–1080. doi:10.1111/j.1471-4159.2004.02927.x

Ishizaki, H., Miyoshi, J., Kamiya, H., Togawa, A., Tanaka, M., Sasaki, T., … Takai, Y. (2000). Role of rab GDP dissociation inhibitor alpha in regulating plasticity of hippocampal neurotransmission. Proc Natl Acad Sci U S A, 97(21), 11587–11592. doi:10.1073/pnas.97.21.11587

Ito, E., Okada, R., Sakamoto, Y., Otshuka, E., Mita, K., Okuta, A., … Sakakibara, M. (2012). Insulin and memory in Lymnaea. Acta Biol Hung, 63 Suppl 2, 194–201. doi:10.1556/ABiol.63.2012.Suppl.2.25

Janse, C., Slob, W., Popelier, C. M., & Vogelaar, J. W. (1988). Survival characteristics of the mollusc Lymnaea stagnalis under constant culture conditions: effects of aging and disease. Mech Ageing Dev, 42(3), 263–274. doi:10.1016/0047-6374(88)90052-8

Jimenez, C. R., ter Maat, A., Pieneman, A., Burlingame, A. L., Smit, A. B., & Li, K. W. (2004). Spatio-temporal dynamics of the egg-laying-inducing peptides during an egg-laying cycle: a semiquantitative matrix-assisted laser desorption/ionization mass spectrometry approach. J Neurochem, 89(4), 865–875. doi:10.1111/j.1471-4159.2004.02353.x

Ju, P., & Cui, D. (2016). The involvement of N-methyl-D-aspartate receptor (NMDAR) subunit NR1 in the pathophysiology of schizophrenia. Acta Biochim Biophys Sin (Shanghai), 48(3), 209–219. doi:10.1093/abbs/gmv135

Kang, H. J., Kawasawa, Y. I., Cheng, F., Zhu, Y., Xu, X., Li, M., … Sestan, N. (2011). Spatio-temporal transcriptome of the human brain. Nature, 478(7370), 483–489. doi:10.1038/nature10523

Kim, D., Langmead, B., & Salzberg, S. L. (2015). HISAT: a fast spliced aligner with low memory requirements. Nat Methods, 12(4), 357–360. doi:10.1038/nmeth.3317

Kojima, S., Nanakamura, H., Nagayama, S., Fujito, Y., & Ito, E. (1997). Enhancement of an inhibitory input to the feeding central pattern generator in Lymnaea stagnalis during conditioned taste-aversion learning. Neurosci Lett, 230(3), 179–182. doi:10.1016/s0304-3940(97)00507-7

Kojima, S., Sunada, H., Mita, K., Sakakibara, M., Lukowiak, K., & Ito, E. (2015). Function of insulin in snail brain in associative learning. J Comp Physiol A Neuroethol Sens Neural Behav Physiol, 201(10), 969–981. doi:10.1007/s00359-015-1032-5

Kuroda, R., & Abe, M. (2020). The pond snail Lymnaea stagnalis. Evodevo, 11(1), 24. doi:10.1186/s13227-020-00169-4

Lambe, E. K., Fillman, S. G., Webster, M. J., & Shannon Weickert, C. (2011). Serotonin receptor expression in human prefrontal cortex: balancing excitation and inhibition across postnatal development. PLoS One, 6(7), e22799. doi:10.1371/journal.pone.0022799

Lansdell, S. J., Collins, T., Goodchild, J., & Millar, N. S. (2012). The Drosophila nicotinic acetylcholine receptor subunits Dalpha5 and Dalpha7 form functional homomeric and heteromeric ion channels. BMC Neurosci, 13, 73. doi:10.1186/1471-2202-13-73

Law, A. J., Weickert, C. S., Webster, M. J., Herman, M. M., Kleinman, J. E., & Harrison, P. J. (2003). Expression of NMDA receptor NR1, NR2A and NR2B subunit mRNAs during development of the human hippocampal formation. Eur J Neurosci, 18(5), 1197–1205. doi:10.1046/j.1460-9568.2003.02850.x

Liao, Y., Smyth, G. K., & Shi, W. (2014). featureCounts: an efficient general purpose program for assigning sequence reads to genomic features. Bioinformatics, 30(7), 923–930. doi:10.1093/bioinformatics/btt656

Liu, M. M., Davey, J. W., Jackson, D. J., Blaxter, M. L., & Davison, A. (2014). A conserved set of maternal genes? Insights from a molluscan transcriptome. Int J Dev Biol, 58(6-8), 501–511. doi:10.1387/ijdb.140121ad

Love, M. I., Huber, W., & Anders, S. (2014). Moderated estimation of fold change and dispersion for RNA-seq data with DESeq2. Genome Biol, 15(12), 550. doi:10.1186/s13059-014-0550-8

Lu, M. R., Lai, C. K., Liao, B. Y., & Tsai, I. J. (2020). Comparative Transcriptomics across Nematode Life Cycles Reveal Gene Expression Conservation and Correlated Evolution in Adjacent Developmental Stages. Genome Biol Evol, 12(7), 1019–1030. doi:10.1093/gbe/evaa110

Maasz, G., Zrinyi, Z., Reglodi, D., Petrovics, D., Rivnyak, A., Kiss, T., … Pirger, Z. (2017). Pituitary adenylate cyclase-activating polypeptide (PACAP) has a neuroprotective function in dopamine-based neurodegeneration in rat and snail parkinsonian models. Dis Model Mech, 10(2), 127–139. doi:10.1242/dmm.027185

Mersman, B. A., Jolly, S. N., Lin, Z., & Xu, F. (2020). Gap Junction Coding Innexin in Lymnaea stagnalis: Sequence Analysis and Characterization in Tissues and the Central Nervous System. Front Synaptic Neurosci, 12, 1. doi:10.3389/fnsyn.2020.00001

Mistry, J., Finn, R. D., Eddy, S. R., Bateman, A., & Punta, M. (2013). Challenges in homology search: HMMER3 and convergent evolution of coiled-coil regions. Nucleic Acids Res, 41(12), e121. doi:10.1093/nar/gkt263

Montellano, O. d. (2015). Cytochrome P450: structure, mechanism and biochemistry (O. d. Montellano Ed. 4th ed.): Kluwer Academic/Plenum Publishers.

Monyer, H., Burnashev, N., Laurie, D. J., Sakmann, B., & Seeburg, P. H. (1994). Developmental and regional expression in the rat brain and functional properties of four NMDA receptors. Neuron, 12(3), 529–540. doi:10.1016/0896-6273(94)90210-0

Moroz, L. L., Edwards, J. R., Puthanveettil, S. V., Kohn, A. B., Ha, T., Heyland, A., … Kandel, E. R. (2006). Neuronal transcriptome of Aplysia: neuronal compartments and circuitry. Cell, 127(7), 1453–1467. doi:10.1016/j.cell.2006.09.052

Moroz, L. L., & Kohn, A. B. (2010). Do different neurons age differently? Direct genome-wide analysis of aging in single identified cholinergic neurons. Front Aging Neurosci, 2. doi:10.3389/neuro.24.006.2010

Moskalev, A. A., Shaposhnikov, M. V., Zemskaya, N. V., Koval Lcapital A, C., Schegoleva, E. V., Guvatova, Z. G., … Kudryavtseva, A. V. (2019). Transcriptome Analysis of Long-lived Drosophila melanogaster E(z) Mutants Sheds Light on the Molecular Mechanisms of Longevity. Sci Rep, 9(1), 9151. doi:10.1038/s41598-019-45714-x

Murakami, J., Okada, R., Sadamoto, H., Kobayashi, S., Mita, K., Sakamoto, Y., … Ito, E. (2013). Involvement of insulin-like peptide in long-term synaptic plasticity and long-term memory of the pond snail Lymnaea stagnalis. J Neurosci, 33(1), 371–383. doi:10.1523/JNEUROSCI.0679-12.2013

Nance, M. A. (2017). Genetics of Huntington disease. Handb Clin Neurol, 144, 3–14. doi:10.1016/B978-0-12-801893-4.00001-8

Nikolac Perkovic, M., & Pivac, N. (2019). Genetic Markers of Alzheimer’s Disease. Adv Exp Med Biol, 1192, 27–52. doi:10.1007/978-981-32-9721-0_3

Onizuka, S., Shiraishi, S., Tamura, R., Yonaha, T., Oda, N., Kawasaki, Y., … Tsuneyoshi, I. (2012). Lidocaine treatment during synapse reformation periods permanently inhibits NGF-induced excitation in an identified reconstructed synapse of Lymnaea stagnalis. J Anesth, 26(1), 45–53. doi:10.1007/s00540-011-1257-6

Ovsepian, S. V. (2017). The birth of the synapse. Brain Struct Funct, 222(8), 3369–3374. doi:10.1007/s00429-017-1459-2

Pacifico, R., MacMullen, C. M., Walkinshaw, E., Zhang, X., & Davis, R. L. (2018). Brain transcriptome changes in the aging Drosophila melanogaster accompany olfactory memory performance deficits. PLoS One, 13(12), e0209405. doi:10.1371/journal.pone.0209405

Papke, R. L., Dwoskin, L. P., & Crooks, P. A. (2007). The pharmacological activity of nicotine and nornicotine on nAChRs subtypes: relevance to nicotine dependence and drug discovery. J Neurochem, 101(1), 160–167. doi:10.1111/j.1471-4159.2006.04355.x

Pereda, A. E. (2014). Electrical synapses and their functional interactions with chemical synapses. Nat Rev Neurosci, 15(4), 250–263. doi:10.1038/nrn3708

Pertea, M., Pertea, G. M., Antonescu, C. M., Chang, T. C., Mendell, J. T., & Salzberg, S. L. (2015). StringTie enables improved reconstruction of a transcriptome from RNA-seq reads. Nat Biotechnol, 33(3), 290–295. doi:10.1038/nbt.3122

Rackauskas, M., Neverauskas, V., & Skeberdis, V. A. (2010). Diversity and properties of connexin gap junction channels. Medicina (Kaunas), 46(1), 1–12.

Sadamoto, H., Takahashi, H., Okada, T., Kenmoku, H., Toyota, M., & Asakawa, Y. (2012). De novo sequencing and transcriptome analysis of the central nervous system of mollusc Lymnaea stagnalis by deep RNA sequencing. PLoS One, 7(8), e42546. doi:10.1371/journal.pone.0042546

Seppala, O., Walser, J. C., Cereghetti, T., Seppala, K., Salo, T., & Adema, C. M. (2021). Transcriptome profiling of Lymnaea stagnalis (Gastropoda) for ecoimmunological research. BMC Genomics, 22(1), 144. doi:10.1186/s12864-021-07428-1

Seshadri, S., Klaus, A., Winkowski, D. E., Kanold, P. O., & Plenz, D. (2018). Altered avalanche dynamics in a developmental NMDAR hypofunction model of cognitive impairment. Transl Psychiatry, 8(1), 3. doi:10.1038/s41398-017-0060-z

Shavlakadze, T., Morris, M., Fang, J., Wang, S. X., Zhu, J., Zhou, W., … Glass, D. J. (2019). Age-Related Gene Expression Signature in Rats Demonstrate Early, Late, and Linear Transcriptional Changes from Multiple Tissues. Cell Rep, 28(12), 3263–3273 e3263. doi:10.1016/j.celrep.2019.08.043

Sodhi, M. S., & Sanders-Bush, E. (2004). Serotonin and brain development. Int Rev Neurobiol, 59, 111–174. doi:10.1016/S0074-7742(04)59006-2

Steen, W. J., Jager, J. H., & Hoven, N. P. V. D. (1968). A Method for Breeding and Studying Freshwater Snails Under Continuous Water Change, With Some Remarks On Growth and Reproduction in Lymnaea Stagnalis (L.). Netherlands Journal of Zoology, 19, 131–139.

Sunada, H., Watanabe, T., Hatakeyama, D., Lee, S., Forest, J., Sakakibara, M., … Lukowiak, K. (2017). Pharmacological effects of cannabinoids on learning and memory in Lymnaea. J Exp Biol, 220(Pt 17), 3026–3038. doi:10.1242/jeb.159038

Swinton, C., Swinton, E., Shymansky, T., Hughes, E., Zhang, J., Rothwell, C., … Lukowiak, K. (2019). Configural learning: a higher form of learning in Lymnaea. J Exp Biol, 222(Pt 3). doi:10.1242/jeb.190405

Syed, N. I., & Winlow, W. (1991). Coordination of locomotor and cardiorespiratory networks of Lymnaea stagnalis by a pair of identified interneurones. J Exp Biol, 158, 37–62.

Tan, R., & Lukowiak, K. (2018). Combining Factors That Individually Enhance Memory in Lymnaea. Biol Bull, 234(1), 37–44. doi:10.1086/697197

Tarkhov, A. E., Alla, R., Ayyadevara, S., Pyatnitskiy, M., Menshikov, L. I., Shmookler Reis, R. J., & Fedichev, P. O. (2019). A universal transcriptomic signature of age reveals the temporal scaling of Caenorhabditis elegans aging trajectories. Sci Rep, 9(1), 7368. doi:10.1038/s41598-019-43075-z

Taylor, B. E., & Lukowiak, K. (2000). The respiratory central pattern generator of Lymnaea: a model, measured and malleable. Respir Physiol, 122(2-3), 197–207. doi:10.1016/s0034-5687(00)00159-6

Team, R. C. (2017). R: A language and environment for statistical computing. Vienna, Austria: R Foundation for Statistical Computing. Retrieved from https://www.r-project.org/

Tebbenkamp, A. T., Willsey, A. J., State, M. W., & Sestan, N. (2014). The developmental transcriptome of the human brain: implications for neurodevelopmental disorders. Curr Opin Neurol, 27(2), 149–156. doi:10.1097/WCO.0000000000000069

Teranishi, Y., Inoue, M., Yamamoto, N. G., Kihara, T., Wiehager, B., Ishikawa, T., … Tjernberg, L. O. (2015). Proton myo-inositol cotransporter is a novel gamma-secretase associated protein that regulates Abeta production without affecting Notch cleavage. FEBS J, 282(17), 3438–3451. doi:10.1111/febs.13353

Teunissen, Y., Geraerts, W. P., van Heerikhuizen, H., Planta, R. J., & Joosse, J. (1992). Molecular cloning of a cDNA encoding a member of a novel cytochrome P450 family in the mollusc Lymnaea stagnalis. J Biochem, 112(2), 249–252. doi:10.1093/oxfordjournals.jbchem.a123885

van Nierop, P., Bertrand, S., Munno, D. W., Gouwenberg, Y., van Minnen, J., Spafford, J. D., … Smit, B. (2006). Identification and functional expression of a family of nicotinic acetylcholine receptor subunits in the central nervous system of the mollusc Lymnaea stagnalis. J Biol Chem, 281(3), 1680–1691. doi:10.1074/jbc.M508571200

van Nierop, P., Keramidas, A., Bertrand, S., van Minnen, J., Gouwenberg, Y., Bertrand, D., & Smit, A. (2005). Identification of molluscan nicotinic acetylcholine receptor (nAChR) subunits involved in formation of cation-and anion-selective nAChRs. J Neurosci, 25(46), 10617–10626. doi:10.1523/JNEUROSCI.2015-05.2005

Vesterlund, L., Jiao, H., Unneberg, P., Hovatta, O., & Kere, J. (2011). The zebrafish transcriptome during early development. BMC Dev Biol, 11, 30. doi:10.1186/1471-213X-11-30

Vorontsov, D. D., Tsyganov, V. V., & Sakharov, D. A. (2004). Phasic coordination between locomotor and respiratory rhythms in Lymnaea. Real behavior and computer simulation. Acta Biol Hung, 55(1-4), 233–237. doi:10.1556/ABiol.55.2004.1-4.28

Wang, B., Wangkahart, E., Secombes, C. J., & Wang, T. (2019). Insights into the Evolution of the Suppressors of Cytokine Signaling (SOCS) Gene Family in Vertebrates. Mol Biol Evol, 36(2), 393–411. doi:10.1093/molbev/msy230

Xia, X., Ding, M., Xuan, J. F., Xing, J. X., Pang, H., Wang, B. J., & Yao, J. (2018). Polymorphisms in the human serotonin receptor 1B (HTR1B) gene are associated with schizophrenia: a case control study. BMC Psychiatry, 18(1), 303. doi:10.1186/s12888-018-1849-x

Xu, Z., Che, T., Li, F., Tian, K., Zhu, Q., Mishra, S. K., … Li, D. (2018). The temporal expression patterns of brain transcriptome during chicken development and ageing. BMC Genomics, 19(1), 917. doi:10.1186/s12864-018-5301-x

Yang, Y. S., & Strittmatter, S. M. (2007). The reticulons: a family of proteins with diverse functions. Genome Biol, 8(12), 234. doi:10.1186/gb-2007-8-12-234

Yeoman, M. S., Kemenes, G., Benjamin, P. R., & Elliott, C. J. (1994). Modulatory role for the serotonergic cerebral giant cells in the feeding system of the snail, Lymnaea. II. Photoinactivation. J Neurophysiol, 72(3), 1372–1382. doi:10.1152/jn.1994.72.3.1372

Zhang, X., Liu, C., Miao, H., Gong, Z. H., & Nordberg, A. (1998). Postnatal changes of nicotinic acetylcholine receptor alpha 2, alpha 3, alpha 4, alpha 7 and beta 2 subunits genes expression in rat brain. Int J Dev Neurosci, 16(6), 507–518. doi:10.1016/s0736-5748(98)00044-6

Zhang, Y., Zhao, J., Zhang, H., Gai, Y., Wang, L., Li, F., … Song, L. (2010). The involvement of suppressors of cytokine signaling 2 (SOCS2) in immune defense responses of Chinese mitten crab Eriocheir sinensis. Dev Comp Immunol, 34(1), 42–48. doi:10.1016/j.dci.2009.08.001

Zoli, M., Pistillo, F., & Gotti, C. (2015). Diversity of native nicotinic receptor subtypes in mammalian brain. Neuropharmacology, 96(Pt B), 302–311. doi:10.1016/j.neuropharm.2014.11.003

