## Supplementary figures and images for "Transcriptome analysis provides genome annotation and expression profiles in the central nervous system of *Lymnaea stagnalis* at different ages"

### Supplemental Figure 1

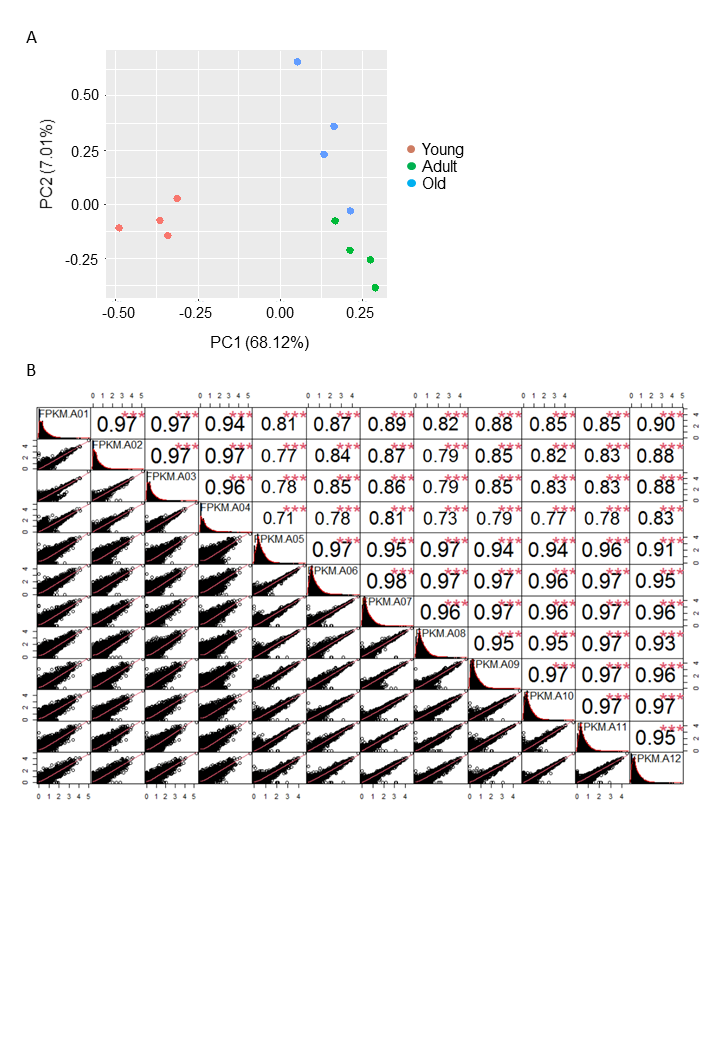

### Supplemental Figure 2

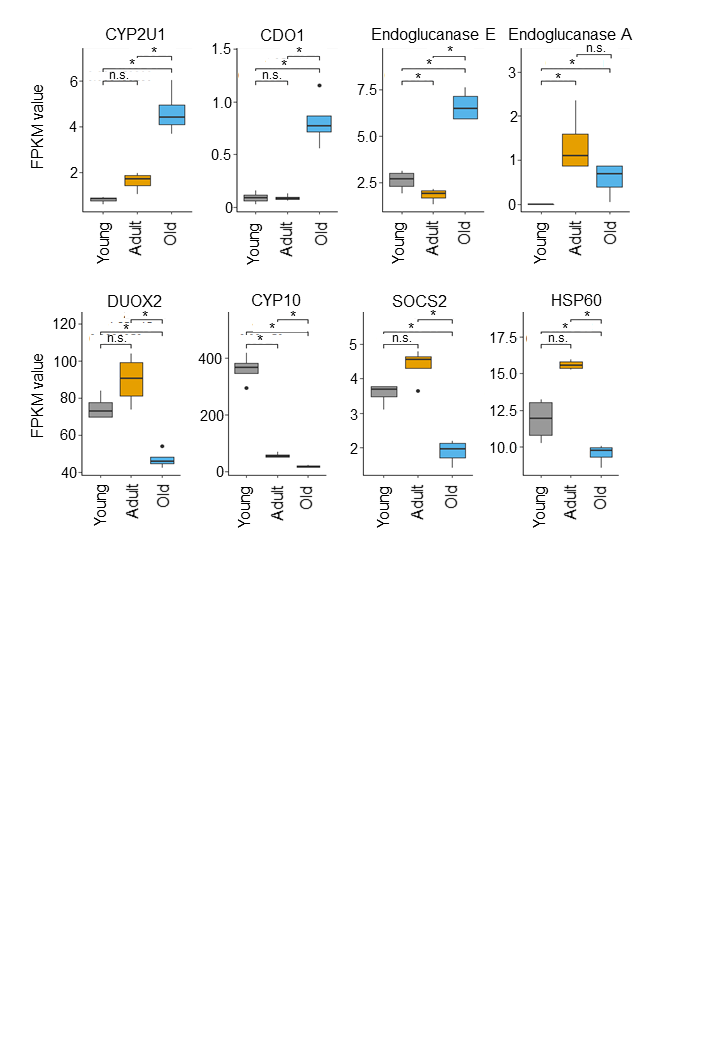

### Supplemental Figure 3

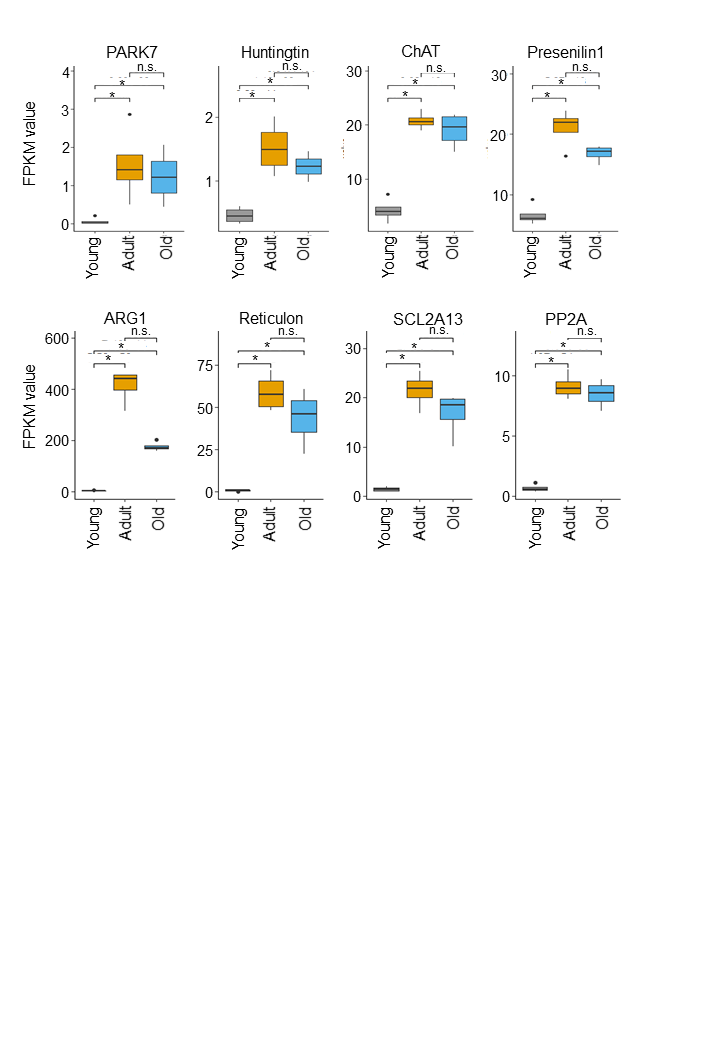
